# Artificial Intelligence-enabled Histological Analysis in Preclinical Respiratory Disease Models: A Scoping Review

**DOI:** 10.1101/2025.10.07.675857

**Authors:** Eva Kuhar, Junsoo Park, Forough Jahandideh, Majid Komeili, Ahmed Sadeknury, Navneet Kang, Prarthna Karunamurthy, Mohammad Reza Zarei, Amir Ebrahimi, Sean E. Gill, Patricia C. Liaw, Dean A. Fergusson, Duncan J. Stewart, Arvind Mer, Manoj M. Lalu, on-behalf-of the LungInsight Collaboration

## Abstract

Histological analysis is a cornerstone of preclinical respiratory disease research, enabling assessment of pathology, therapeutic effects, and mechanisms. However, conventional approaches rely on manual scoring, which is subjective, time-consuming, and difficult to scale due to low throughput and inter-observer variability. Artificial intelligence (AI), particularly deep learning, offers potential to automate histology workflows, but its use and evaluation in preclinical respiratory models have not been synthesized.

We conducted a scoping review following Joanna Briggs Institute guidelines, searching MEDLINE and Embase (inception–January 2025) for preclinical studies using AI to analyze histology in respiratory disease models. Screening, full-text review, and data extraction were performed in duplicate.

Of 6271 studies screened, 29 met inclusion criteria. Most used murine models (76%) and investigated lung cancer (28%), pulmonary fibrosis (24%), or tuberculosis (17%). Hematoxylin and eosin was the most common stain (48%), with others targeting collagen or immune markers. AI tasks included image classification (n=20), segmentation (n=10), and object detection (n=4), predominantly using convolutional neural networks (69%). Preprocessing methods (e.g., stain normalization) were common, but annotation and training practices were inconsistently reported. Performance was generally high (accuracy ≥90%; 7 studies) though validation metrics varied, and external validation was absent. Most studies used “black box” models, with minimal application of explainability techniques. Reproducibility measures, such as sharing datasets or code were rarely reported.

AI tools are poised to transform histological analysis in preclinical respiratory research. By addressing gaps in validation, transparency, and standardization, the field can harness these technologies to deliver robust, efficient, and scalable workflows.

Registration: Open Science Framework https://doi.org/10.17605/OSF.IO/NM94E

## INTRODUCTION

In preclinical respiratory disease research, histological analysis serves as a cornerstone, providing essential insights into structural abnormalities, inflammation, and fibrosis. (1) However, scoring histological features in preclinical animal models remains a significant challenge. Traditional manual assessments are time-intensive, laborious, and prone to inter-subject variability, hindering efficiency and reproducibility in preclinical studies. (2–4) These limitations create difficulties in both the evaluation of experimental therapies and the translation of preclinical findings into clinical applications.

Recent studies have shown that artificial intelligence/machine learning (AI/ML) technologies are emerging as tools to automate and enhance histological analysis in respiratory diseases. (5–7) AI-powered algorithms, particularly deep learning approaches, have the potential to enable rapid, objective, and high-throughput quantification of histological patterns. In preclinical research, AI, and particularly convolutional neural networks (CNNs), have shown promise in automating histological scoring of fibrotic lung disease in mice (8), detecting tumorous lesions on whole-slide images (9), and classifying tumor phenotypes in experimental lung cancer studies. (10) These advancements could enhance the reproducibility and efficiency of histological scoring, reducing/eliminating the inherent variability in manual assessment. Despite these promising developments, there are still underlying challenges to be addressed, such as the need for large, well-annotated histology datasets, validation across distinct animal models of lung injury, and the standardization of AI-based scoring methods. Moreover, the use of AI approaches in respiratory disease research is a rapidly evolving field, with new technologies and methodologies being developed at an accelerating pace. Therefore, a synthesis of AI technology and machine learning in animal models of respiratory disease is needed to inform the next steps in the development and implementation of these technologies for histological analysis.

This scoping review synthesizes current applications of AI-enabled histological analysis in preclinical respiratory disease models. We focus on preclinical studies since animal models allow controlled experimental conditions, precise timing of sample collection, and the ability to obtain histological data across multiple disease stages or treatment time points - approaches that are rarely feasible in clinical research, where tissue access is often opportunistic (e.g., during surgery, biopsy, or at autopsy). Moreover, histological analysis is a gold standard outcome measure in many preclinical respiratory disease models (11, 12), providing direct assessment of disease pathogenesis, therapeutic efficacy, and mechanistic insights. By evaluating current methodologies, we can help identify technical advantages, limitations, and critical knowledge gaps to advance robust AI tools for preclinical respiratory research. Addressing these challenges is necessary for accelerating the translational gap and ensuring that AI-driven histological insights contribute meaningfully to preclinical research.

## METHODS

### Protocol and registration

We conducted this scoping review in accordance with the Joanna Briggs Institute (JBI) methodological guidelines (Peters, 2022) and with the methodological framework initially proposed by Arksey and O’Malley (2005) and later updated by Levac et al (2010). (13–15) This review is reported in accordance with the Preferred Reporting Items for Systematic Reviews and Meta-Analyses extension for scoping reviews (PRISMA-ScR) checklist (a completed checklist can be found in Appendix 1). The protocol can be found in Open Science Framework (https://doi.org/10.17605/OSF.IO/NM94E).

### Identifying the research question

Our research question was: “What AI/ML tools exist in assessing histological features in respiratory diseases that can be used in animal models of ALI?” We aimed to comprehensively synthesize preclinical studies using AI/ML technologies for lung histology in animal models of respiratory disease.

### Information sources and literature search

We developed a systematic search strategy (Appendix 2) in collaboration with an information specialist (Risa Shorr, MLS, The Ottawa Hospital Learning Services). Our search strategy was applied to PubMed (MEDLINE) and Embase (search last updated on January 20^th^, 2025), and no date restrictions were applied to be as comprehensive as possible. Keywords used in our search included “respiratory tract disease”, “artificial intelligence”, and other relevant terms related to the disease, methodology and technology used in machine learning. Due to feasibility, we restricted inclusion to peer-reviewed manuscripts of preclinical, primary studies written in English.

### Eligibility criteria

We included experimental studies investigating the use of an AI/ML tool to assess histological evidence of lung pathology in animal models of human respiratory diseases. To ensure clarity in study selection, AI was defined as the study of intelligent systems that perform tasks typically requiring human intelligence. (16) Machine learning (ML), a subfield of AI, refers to approaches in which intelligence emerges from learning patterns in annotated data. (17) We included all species and preclinical respiratory disease models. Algorithmic methods were included if they demonstrated sufficient learning or reasoning capabilities (i.e., characteristics of AI). For this review, pathohistological features were defined as histological structures that differentiate between healthy and diseased lung tissue. Studies were excluded if they (i) did not use AI/ML specifically for histopathological assessment, (ii) primarily focused on non-histological imaging modalities (e.g., x-rays, computed tomography, magnetic resonance imaging), or (iii) used histopathological analysis solely as a validation tool for another primary assessment method.

### Study screening and selection

All citations retrieved through the search were uploaded into DistillerSR (Evidence Partners, Ottawa), a cloud-based, audit-ready software that streamlines scoping review workflows, including title and abstract screening, full-text review, and duplicate removal. Two independent reviewers screened articles in duplicate by titles/abstracts and then by full text. Discrepancies between the reviewers were resolved through discussion or by a senior author, if needed.

### Data extraction and synthesis

A standardized data extraction form was developed to systematically collect information on study characteristics, methodologies, AI-based tools, and results. Two independent reviewers extracted data in duplicate, and discrepancies were resolved through discussion or by a senior author, if needed. The extracted information encompassed five major domains:

- *Study characteristics and animal model data*: This domain included details on the study’s publication year and author information. It also captured key characteristics of the animal model, such as the species, sex, weight, age, and the specific respiratory disease being modelled. Additionally, the method of disease induction and the histological features assessed were extracted.
- *Scoring and imaging data*: Information was collected on the imaging and histological assessment methods used in each study. This included the type of microscope used, the method of image acquisition, and the magnification applied for analysis. The staining technique, histological scoring system, and the number of training images used in AI model development were also collected.
- *AI-based tool characteristics*: Information captured included the name and type of automation system or technology (e.g., convolutional neural network [CNN], support vector machine [SVM], image analysis platforms such as QuPath or HALO), the level of AI autonomy (e.g., decision-support systems where humans make the final judgment, partially automated systems that perform tasks with limited human oversight, or fully automated systems operating independently) used for developing the system, specific AI techniques and tasks (e.g., image classification, segmentation), and any interpretability considerations.
- *AI model performance and robustness*: Data were collected on the performance of the AI model, including accuracy, and other performance metrics, as well as external validation. Assessments of inter- and intra-observer variability and measures of agreement reliability were also recorded. The accessibility of the AI software, such as whether the code was publicly shared, was also noted.
- *Outcomes and challenges*: The comparison of AI-generated histological feature scores with human expert assessments, challenges/limitations, and the study authors’ conclusions regarding the applicability of the AI tool were also collected.

### Data analysis and reporting

Extracted data were synthesized using descriptive statistics and qualitative synthesis. Study and animal model characteristics and categorical data were summarized in tabular format. For AI model performance, reported accuracy metrics were compared with human expert assessments where available. Qualitative findings, including study conclusions and limitations, were analyzed to identify common patterns across studies. Figures and tables were generated to present key findings. All analyses were conducted in R (version 4.4.3, Windows UCRT build) using the packages tidyverse (2.0.0), readxl (1.4.5), circlize (0.4.16), and png (0.1.8).

## RESULTS

### Overview of included studies

Our systematic search identified 7,806 records, with 6,271 unique records screened, and 29 studies meeting the inclusion criteria for this review (Supplementary Figure S1). These studies were published between 2008 and 2025. 59% (n=17) of studies were published in the last five years.

The most frequently investigated respiratory conditions were lung cancer (n=8, 28%), pulmonary fibrosis (n=7, 24%), and tuberculosis (n=5, 17%) (Table 1). Pulmonary hemorrhage (10%, n=3) and Acute Lung Injury (ALI) models were also investigated, with the latter specifically addressing SARS-CoV-2 (n=2) and bacterial pneumonia (n=1). Other conditions studied included chronic obstructive pulmonary disease (COPD; n=2, 7%), and bronchopulmonary dysplasia (BPD; n=1, 3%, Table 1, Figure 1A).

**Figure 1.**
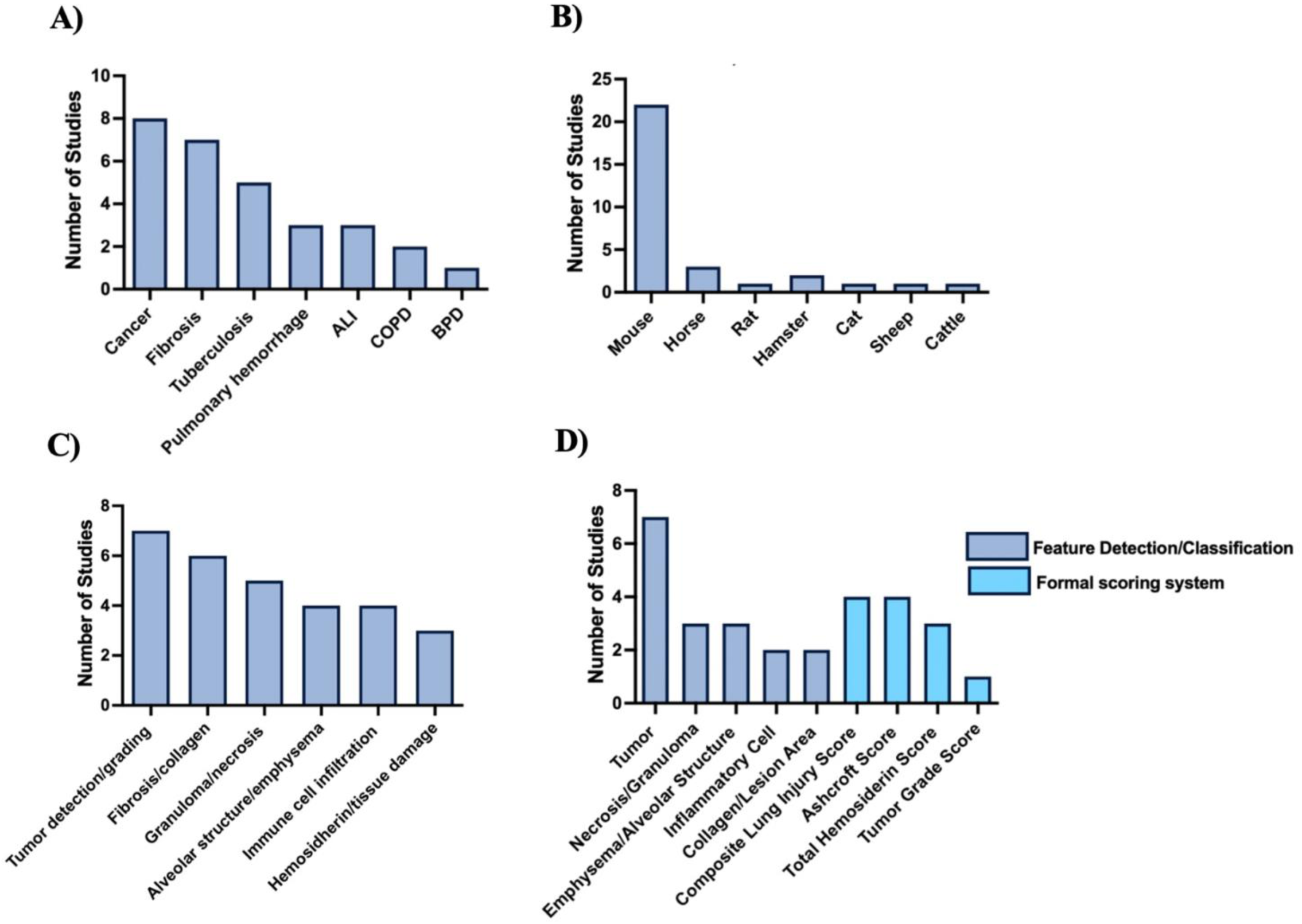
Respiratory disease models, animal species, histological features, and scoring systems investigated in included studies. Frequency of (A) respiratory disease models studied, (B) animal species used, (C) histological features evaluated in artificial intelligence-driven histology research, (D) use of feature detection/classification versus formal scoring systems across studies. Abbreviations: ALI, acute lung injury; COPD, chronic obstructive pulmonary disease; BPD, bronchopulmonary dysplasia.

**Table 1.** Characteristics of studies applying artificial intelligence approaches to respiratory disease histology. Abbreviations: AFB, acid-fast bacilli; BPD, bronchopulmonary dysplasia; COPD, chronic obstructive pulmonary disease; DAD, diffuse alveolar damage; DO, diversity outbred; EIPH, exercise-induced pulmonary hemorrhage; ET, endotracheal; F, female; GEMMs, genetically engineered mouse models; IN, intranasal; IP, intraperitoneal; IT, intratracheal; IV, intravenous; LRT, lower respiratory tract; M, male; NS, not specified; NSCLC, non-small cell lung cancer; O₂, oxygen; SARS-CoV-2, severe acute respiratory syndrome coronavirus 2. SD, Sprague Dawley.

| First author | Year | Species | Strain | Age (weeks) | Sex | Method of disease induction | Respiratory disease modeled | Histological features scored |
| --- | --- | --- | --- | --- | --- | --- | --- | --- |
| Maglogiannis, I. | 2008 | Mouse | NS | 6–8 | NS | IV bleomycin | Fibrosis | Interstitial/alveolar inflammation |
| Marcos, J. | 2015 | Mouse | A/J | 11 | M | IT porcine pancreatic elastase | Emphysema COPD | Emphysema severity, airspace opacities |
| Jule, Y. | 2017 | Mouse | C57/BL6 | 8-12 | M | IT bleomycin | Fibrosis | Pulmonary tissue density |
| Dietert, K. | 2018 | Mouse | C57BL/6 | 8-10 | F | IN *S. pneumoniae* + influenza A | Pneumonia, Influenza A | CD68, neutrophil elastase, CD3, CD45R |
| Heinemann, F. | 2018 | Mouse | C57BL/6J, C57BL/6N | 8-12 + 18-20 mo | Both | IT Bleomycin | Lung fibrosis | Fibrotic masses, immune cell infiltration |
| Seger, S. | 2018 | Rat | Wistar, SD | NS | M | IT bleomycin | IPF | Fibrotic masses, alveolar wall collagen |
| Beamer, G. | 2019 | Mouse | DO | 8 | F | IN *M. tuberculosis* (Erdman) | Tuberculosis | Granuloma necrosis |
| Bug, D. | 2019 | Mouse | NS | NS | NS | NS | NSCLC | Tumor/stromal area |
| Ben-Hur, A. | 2020 | Mouse | C3HeB/FeJ | 6-8 | F | Inhaled *M. tuberculosis* | Tuberculosis | Necrotic granuloma, neutrophil infiltration |
| Niazi, M. | 2020 | Mouse | DO, C57BL/6J | 8-10 | F | IN *M. tuberculosis* (Erdman) | Tuberculosis | Granuloma necrosis |
| Marzahl, C | 2020 | Horse | NS | NS | NS | Clinical LRT symptoms | EIPH | Hemosiderin Count |
| Bouzin, C. | 2020 | Mouse | C57BL/6J | 8 | M | ET bleomycin | Fibrosis | Collagen deposits, fiber segmentation |
| Harmon, S. | 2021 | Mouse | GEMMs | NS | Both | Adenovirus ‘Cre’ | Adenocarcinoma | Tumorous lesions, tumour burden |
| Salsabili, S. | 2021 | Mouse | NS | NS | NS | High O<sub>2</sub> + LPS | BPD | Septal crossing intersection |
| Kang, J. | 2022 | Mouse | C57BL/6 | NS | F | IV (B16-F10) melanoma cells | Metastatic lung cancer | Metastatic tumors, tumorous lesions |
| Marzahl, C. | 2022 | Horse, Cat | NS | NS | NS | NS | Pulmonary hemosiderophages | Hemosiderin count |
| Martinot, A. | 2022 | Hamster | Golden Syrian | 9 | Both | IN SARS-CoV-2 | SARS-CoV-2 | Airways, interstitium, vasculature, alveolar spaces, edema, DAD |
| Bartel, A. | 2023 | Horse | NS | NS | NS | Routine asthma evaluation | EIPH | Hemosiderin Count |
| Ducote, T. | 2023 | Mouse | GEMMs | NS | NS | Inhaled Cre/FlpO virus | Lung cancer | Tumor microenvironment, immune infiltration |
| Palmarini, M. | 2023 | Hamster | Syrian | 8-12 | Both | IN SARS-CoV-2 | SARS-CoV-2 | Alveolar hyperplasia, macrophage infiltration, immune response |
| Ram, S. | 2023 | Mouse | K-rasG12D | NS | NS | Genetically engineered | Lung cancer | Tumorous lesions |
| Chandra, D. | 2023 | Mouse | C57/BL6, BALB/c | 10 | F | 6-month cigarette smoke exposure | COPD | Chord length (alveolar size) |
| Flores, E. | 2023 | Mouse | C57/BL6 | 8-10 | Both | IT adenovirus-Cre | Lung cancer | Tumor grading, cellular differentiation |
| Murgo, A. | 2023 | Mouse | C57/BL/6J | 6-7 | M | Oropharyngeal Bleomycin | Fibrosis | Ashcroft score, fibrotic changes |
| Vincenzi, E. | 2024 | Mouse | C57/BL6 | 8-10 | M | Oropharyngeal Bleomycin | Fibrosis | Ashcroft score, collagen deposition |
| Liu, S. | 2024 | Sheep | NS | NS | NS | Ovine pulmonary adenocarcinoma | Lung cancer | Tumorous lesions |
| Alnafisah, K. | 2024 | Mouse | BALB/c | >48 | F | IP urethane | Lung cancer | Collagen, fibrillar alterations |
| Koyuncu, D. | 2024 | Mouse | DO | 8-10 | F | Aerosolized *M. tuberculosis* | Tuberculosis | Perivascular/peribronchiolar granuloma, B-cell infiltration |
| Acharya, V. | 2024 | Mouse | DO | 8 | F | IN *M. tuberculosis* | Tuberculosis | AFB, macrophage infiltration, granulomas |

Murine models were predominantly used (n=22, 76%, Table 1), with commonly cited strains including C57BL/6, BALB/c, C3HeB/FeJ, diversity outbred (DO) mice, and A/J. Equine models comprised 10% of the studies (n=3), all of which investigated pulmonary hemorrhage. Individual studies also used hamster (SARS-CoV-2), sheep (lung cancer), and rat (pulmonary fibrosis) models (each 3%, Table 1, Figure 1B).

### Staining and imaging modalities

Hematoxylin and eosin were the most frequently used staining method (n=14, 48%, Supplementary Table S1, S2), while the remaining studies (n=15, 52%) employed stains tailored to particular histologic features. Collagen was assessed by Masson’s trichrome (n=5), picrosirius red (n=1), and second harmonic generation microscopy (a stain-free optical imaging method, n=1). Prussian blue and modified Turnbull’s blue were used in three studies to quantify hemosiderin as a surrogate for pulmonary hemorrhage. Other specialized staining approaches included RNA in situ hybridization (n=2) targeting viral RNA and immune cell markers (such as TTF1, IBA1, MX1), modified Ziehl-Neelsen (carbolfuchsin, n=1) for *Mycobacterium tuberculosis*, and modified Gill’s stain (n=1) for measuring chord length.

Digitization of stained slides was performed using high-resolution, whole-slide imaging systems, with scanners such as Aperio ScanScope (Leica), NanoZoomer (Hamamatsu), and Zeiss AxioScan frequently cited (Supplementary Table S1). Image acquisition parameters were chosen based on downstream analytical needs. Lower magnifications (≤10×) provided broader architectural context, while higher magnifications (20× and 40×) were most often used for the complex classification and segmentation required for AI modelling. Seven studies utilized ultra-high magnification (≥200×), especially for tasks needing high cellular resolution, such as immunophenotyping. (18–20)

### Histological features and scoring systems

The histological features targeted for AI-based evaluation varied according to the respiratory disease model and the primary histological focus (Figure 1C, Supplementary Table S2). The most common applications involved quantifying fibrotic lesions and collagen deposition (n=8, 28%), followed by tumor detection or grading (n=6, 21%), alveolar architectural changes and emphysema severity (n=4, 14%), granuloma formation and necrosis in tuberculosis (n=4, 14%), immune cell infiltration (n=3, 10%), and hemosiderin accumulation or tissue damage metrics (n=2, 7%).

A variety of histological scoring strategies were used to validate AI outputs. The majority of studies (n=17, 59%) relied on direct feature detection or classification rather than established formal scoring systems (Figure 1D). The majority of tumor-focused studies (n=5) used feature detection approaches, with formal grading systems applied in one study. Similar trends were evident for emphysema, alveolar structure, inflammatory cell assessment, and collagen or lung structure quantification, where studies more often employed AI-based feature extraction over traditional scoring frameworks. Among studies using formal scoring systems, the Ashcroft scale for fibrosis severity and composite lung injury scores were most common (n=4 each), followed by the Total Hemosiderin Score in pulmonary hemorrhage studies (n=3).

### AI methods

#### Image processing workflows

Most studies (83%, n=24) implemented at least one standardized image preprocessing strategy to reduce variability from staining, illumination, and imaging differences. Common steps included artifact removal and exclusion of non-diagnostic tissue regions to ensure analysis focused on biologically relevant areas (n=22). Dataset augmentation through geometric or color transformations was used in 19 (66%) studies to increase sample diversity and enhance model robustness. Stain normalization methods (e.g. Macenko algorithm) were applied in 13 studies (35%) to minimize color inconsistencies from sample preparation and scanning variations. Image contrast enhancement was also used in 13 studies to improve the delineation of histological features and boundary identification. Patch-based tiling was implemented in 13 studies. This approach, which divides whole slides into smaller tiles, enabled efficient gigapixel image processing. Patch sizes and overlap parameters were typically optimized based on model architectures and resolution requirements.

Technical parameters such as section thickness and scanner resolution, reported in seven studies, influenced tissue appearance and downstream analytical performance. Variations in section thickness complicated image registration, particularly when aligning histological slides with complementary modalities like micro-computed tomography. (21)

Four studies used classical image processing pipelines without modern AI, relying on manually engineered features or threshold-based rules. Additionally, two studies implemented hybrid workflows combining traditional processing with machine learning through manual or semi-automated region selection. (22, 23)

#### AI Task Applications

Across the 29 included studies, we identified 34 distinct AI task applications, with several studies implementing multiple task types. The most common applications were image classification (n=20), image segmentation (n=10), and object detection (n=4) (Figure 2, Supplemental Figure S2). Image classification was broadly adopted across disease models, including fibrosis (n=6), cancer (n=5), tuberculosis (n=4), SARS-CoV-2 (n=1), pneumonia (n=1), BPD (n=1), pulmonary hemorrhage (n=1), and COPD (n=1), underscoring its broad applicability (Figure 2). Image segmentation (automated partitioning of images into distinct anatomical regions) was applied in cancer (n=3), fibrosis (n=2), COPD (n=2), SARS-CoV-2 (n=1), tuberculosis (n=1), and BPD (n=1), enabling more precise measurement of either tissue structure, lesion boundaries, or cellular infiltration. Object detection (automated identification and localization of discrete features) was primarily used for pulmonary hemorrhage (n=3), for identifying hemosiderin-containing macrophages in lung sections, and in one pneumonia study that quantified neutrophils and total counts of cells per area.

**Figure 2.**
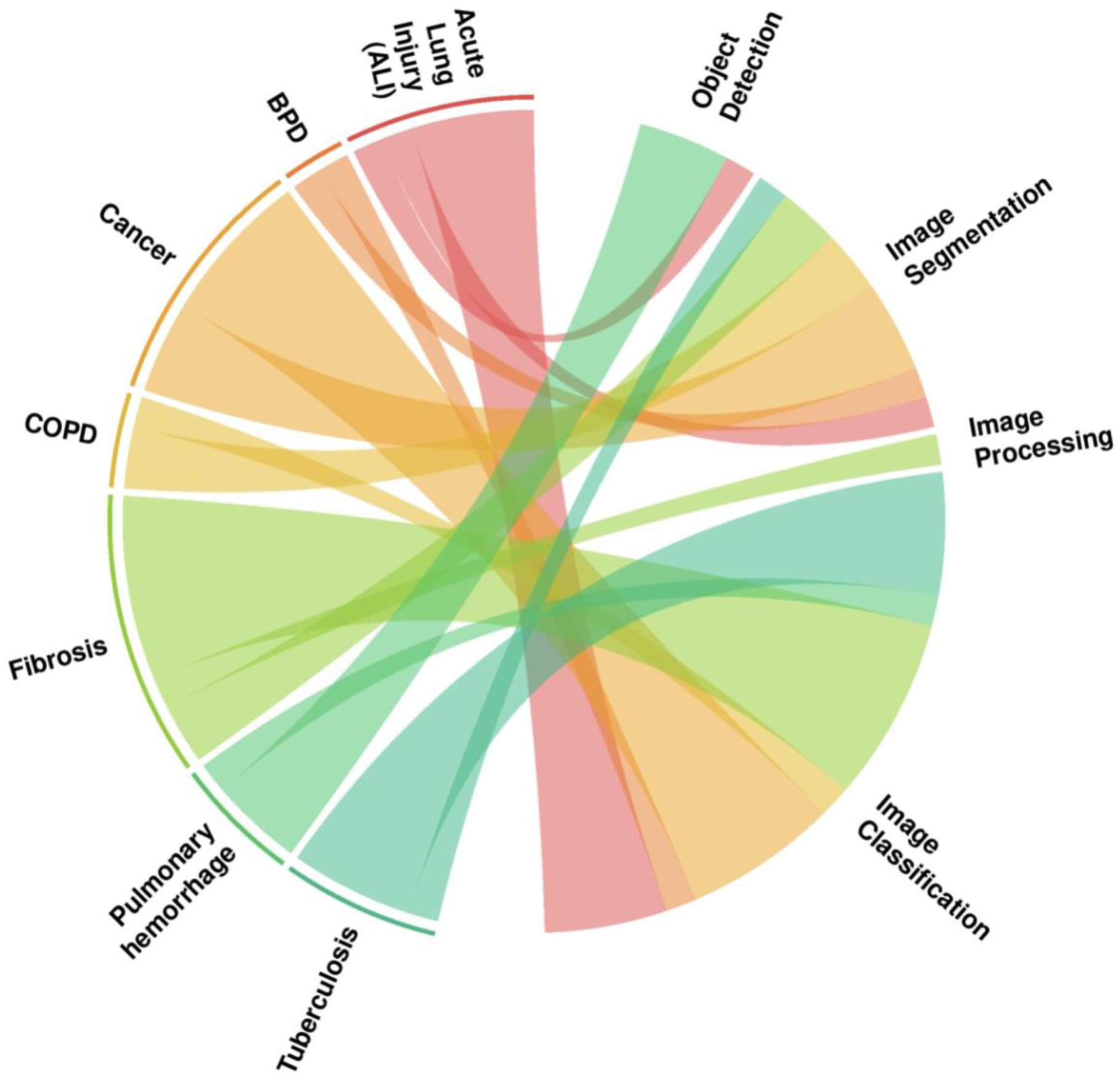
Characteristics of artificial intelligence techniques and their application to preclinical respiratory disease histology. Chord diagram illustrating how different AI techniques (right) are applied across various respiratory diseases (left) in histological studies. Line thickness indicates the number of studies reporting a given pairing.

#### AI technologies, learning approaches, and implementation

The 29 included studies used diverse AI approaches, which were categorized by learning mechanism (i.e. data-driven versus not) and model architecture. Data-driven models infer statistical relationships between pixel values and labels through training, while rule-based methods apply predefined image-processing rules without model training.

Most studies (n=27, 93%) employed data-driven learning approaches, primarily deep learning (n=20, 69%), which learn hierarchical features directly from raw images, often reducing the need for handcrafted feature engineering. CNNs were used for image classification and feature extraction, with specialized architectures (e.g. Inception V3) (24) used to distinguish tissue regions such as tumors, necrosis, or fibrosis. U-Net (25) and DeepLabV3+ (26) were employed for pixel-level segmentation tasks that enabled identification of tumors, necrosis, and fibrosis (Supplementary Table S2). (27, 28) Some incorporated morphometric analysis following AI-driven segmentation, such as U-Net–based exclusion of non-alveolar structures for chord length measurement. (29)

Object detection, such as RetinaNet (7%, n=2), extended beyond simple classification by automating localization and quantification of features (e.g. metastatic foci, hemosiderin-laden macrophages). Attention-based models (including multiple instance learning frameworks) were used in two studies for whole slide analysis, enabling the identification of regions that contribute most significantly to a diagnostic label. Other neural networks included radial basis function models (n=4, 14%) and Bayesian models (n=2, 7%) for tasks where pixel-level detail labels were unavailable.

Traditional machine learning methods were applied in seven studies. This requires domain experts to define specific measurable features (e.g. cell morphometry, texture descriptors, color) from histological images, after which algorithms learn to make diagnostic decisions based on these pre-engineered features. In one study, support vector machines were integrated into multi-step analysis workflows (e.g. initial segmentation of granulomas followed by support vector machine-assisted classification of necrotic regions. (22)

Two studies (7%) implemented rule-based algorithms without any data-driven learning component that relied on manually programmed instructions for pixel-level classification or feature extraction,

Implementation strategies varied significantly across studies. Several studies (31%, n=9) leveraged existing tools for feature extraction, segmentation, or classification rather than developing novel algorithms (e.g. Orbit Image Analysis, Lesion Image Recognition and Analysis (LIRA), HALO). Conversely, a subset of studies (10%, n=3) described building custom multi-stage AI pipelines that combined pre-trained models and classical image processing in a modular fashion, tailoring the methodological approach to the specific complexity of lung histopathology.

#### Annotation practices and slide volume

Annotation methodologies varied substantially across the 29 included studies (Supplementary Table S2). Manual annotation was most common (n=18, 62%) for creating pixel-level segmentations, bounding boxes, or tissue/region-level labels. Twelve of these relied on a single annotator, most commonly a pathologist (n=7), followed by experienced personnel or PhD students (n=3), while two studies did not report qualifications. The remaining six studies used multiple annotators, often with varying expertise, to assess inter-annotator variability.

Semi-automated annotation with human validation was employed in 6 studies (21%). Three studies (10%) used fully automated systems or synthetic data without manual labelling; in two of these, pathologists selected relevant features for the model input but did not annotate the slides.

Most studies (n=22, 76%) reported the number of annotated whole-slide images, which is a proxy for dataset diversity, ranging from 10 to 464 (median 69). Reporting of annotated regions (often stated as ‘tiles ‘or ‘patches’) indicating the extent of model training data was inconsistent. Two studies provided exact tile counts in the hundreds (e.g., 324 tiles from 6 mice), and three in the tens of thousands (e.g., 35,115 tiles). Four studies used less precise descriptors such as ‘thousands of micro-tiles’ or ‘multiple fields per whole slide image’, and the majority of studies (n=20) did not specify tile or patch counts at all. This variability limits direct comparison between datasets and may hinder assessments of model generalizability and reproducibility.

#### Human integration and explainability

Seven studies demonstrated structured integration of AI in human workflows (Supplementary Table S4). One implemented a formal human-in-the-loop approach, while six reported human involvement without operational details. Among the included studies, only one implemented an explainability framework to support the interpretation of the AI model, highlighting that explainability has been overlooked.

#### Validation strategies

Validation approaches varied, with most studies employing internal validation strategies (n=24, 83%). These included dividing available slides into separate sets for model training and testing, or using cross-validation, where models are evaluated on different subsets of the same dataset in rotation. Notably, no studies reported using external validation with fully independent datasets.

#### Overall performance

AI models demonstrated promising performance across a diverse range of tasks (Table 2), with 79% (n=23) reporting at least one validation metric. Validation approaches were heterogeneous (Table 2, Figure 3) where most studies reported a single outcome only (n=20, 69%; e.g. accuracy for classification or Dice similarity for segmentation), while omitting complementary measures (e.g., sensitivity, specificity, confusion matrices) or more comprehensive metrics (e.g. F1 score, area under the curve). Performance varied by task complexity. Simple tasks such as binary classification (e.g., distinguishing tumor-positive and negative regions) achieved the highest accuracy (95 to 99%) and correlation coefficients (up to 0.98), particularly when models were trained with supervised learning on well-annotated datasets. In contrast, more complex tasks such as multilabel classification, spatially resolved cell population analysis, and cross-species prediction showed lower and more variable performance (73 to 75% accuracy), reflecting challenges posed by morphological overlap, cellular heterogeneity, and interspecies variability in preclinical histology.

**Figure 3.**
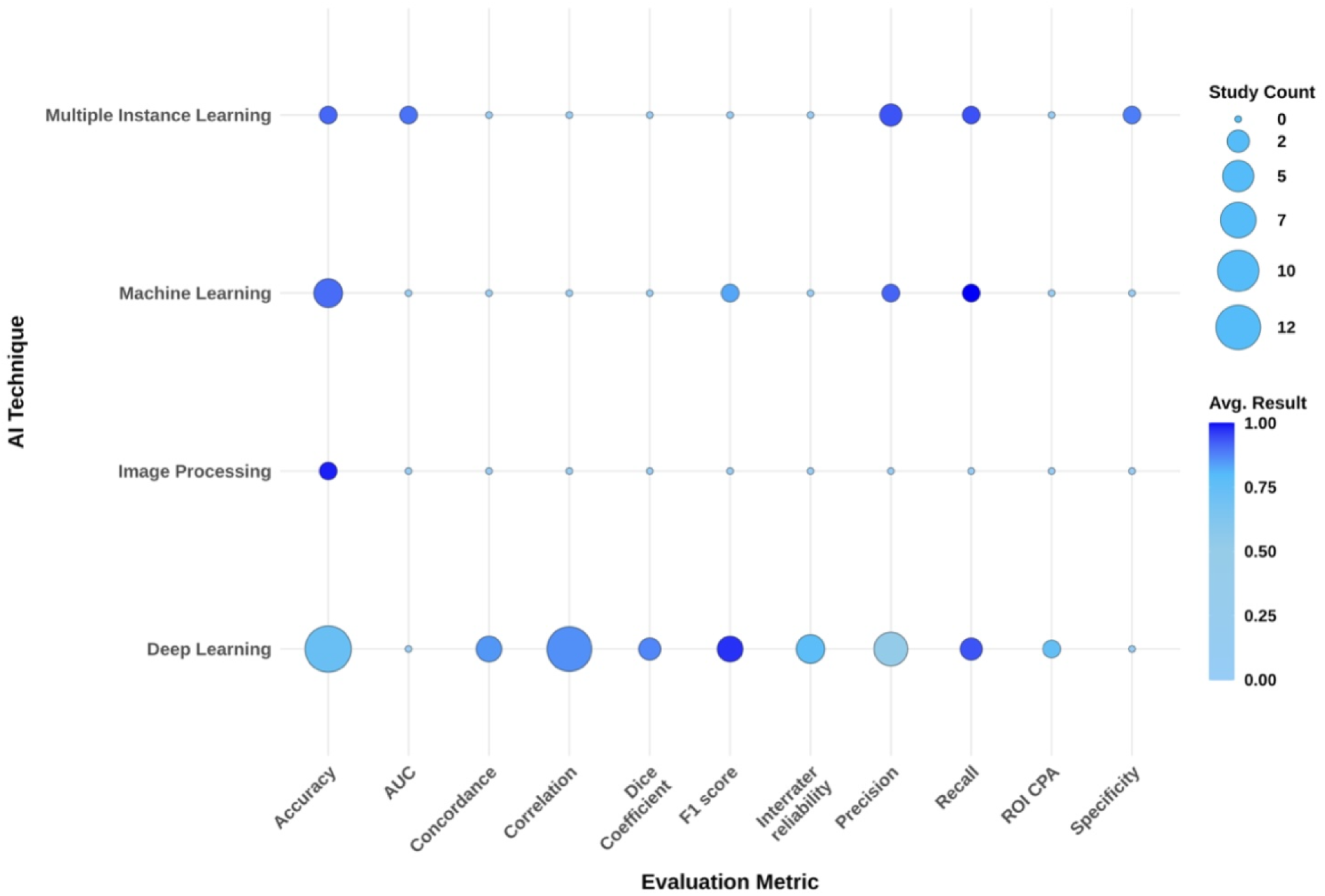
Performance evaluation of artificial intelligence techniques applied in included studies. This dot plot illustrates the distribution and frequency of evaluation metrics used to assess different artificial intelligence techniques across included studies. The size of each dot reflects the number of studies reporting a given metric, while the color intensity indicates the average reported performance. Abbreviations: AI, artificial intelligence; ML, machine learning; AUC, area under the curve; ROI CPA, region of interest classification or prediction accuracy.

**Table 2.**
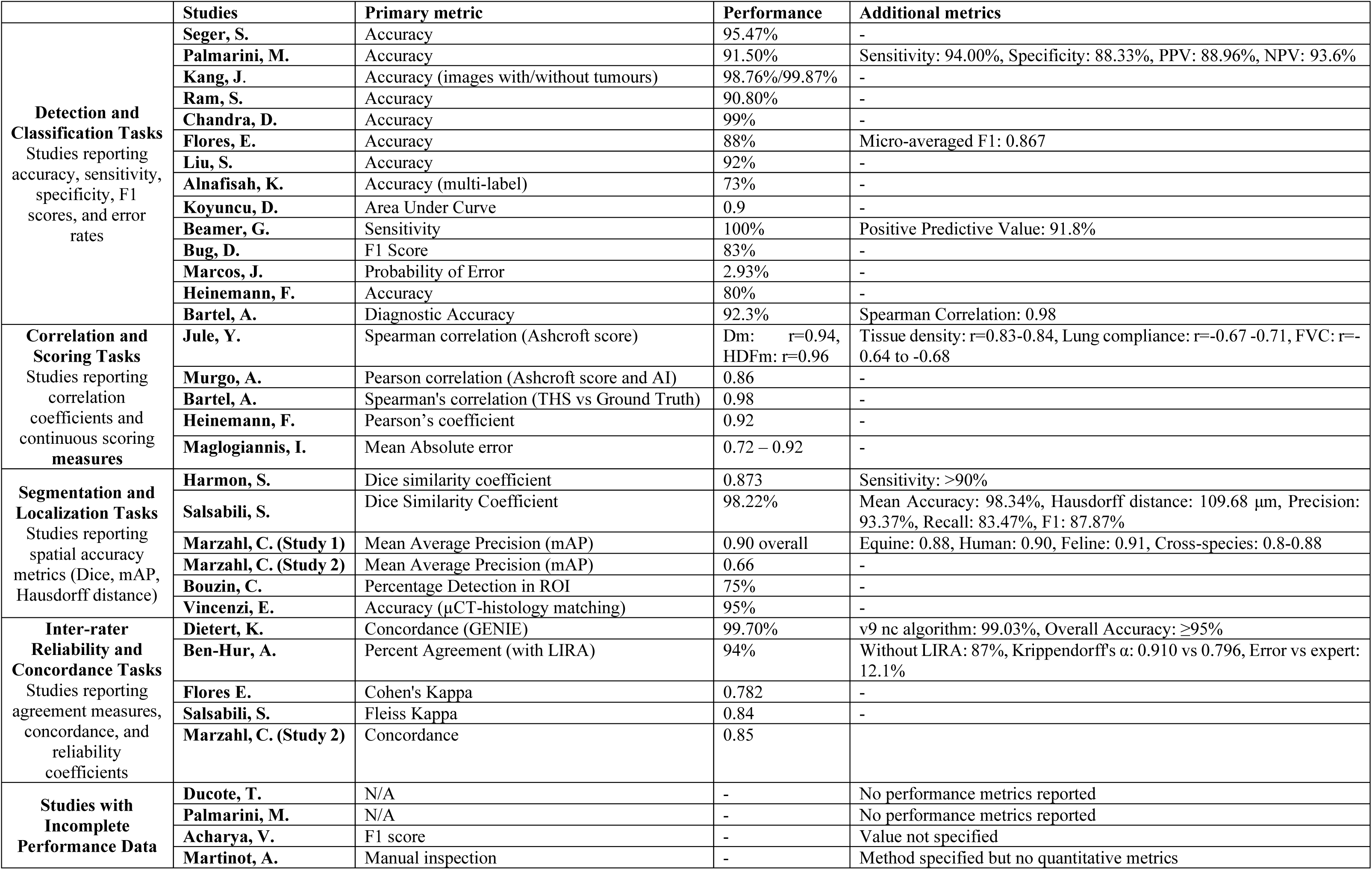
Performance metrics and validity measures of AI models across studies. Abbreviations: PPV, positive prediction value; NPV, negative prediction value; mAP, mean average precision; ROI, region of interest; μCT, micro-computed tomography; Dm, Spearman correlation coefficient (between datasets or scores); FVC, forced vital capacity; GENIE, Genomic Evidence Neoplasia Information Exchange; LIRA, inter-rater agreement; THS, total hemosiderin score.

|  | Studies | Primary metric | Performance | Additional metrics |
| --- | --- | --- | --- | --- |
| **Detection and Classification Tasks**<br>Studies reporting accuracy, sensitivity, specificity, F1 scores, and error rates | Seger, S. | Accuracy | 95.47% | - |
|  | Palmarini, M. | Accuracy | 91.50% | Sensitivity: 94.00%, Specificity: 88.33%, PPV: 88.96%, NPV: 93.6% |
|  | Kang, J. | Accuracy (images with/without tumours) | 98.76%/99.87% | - |
|  | Ram, S. | Accuracy | 90.80% | - |
|  | Chandra, D. | Accuracy | 99% | - |
|  | Flores, E. | Accuracy | 88% | Micro-averaged F1: 0.867 |
|  | Liu, S. | Accuracy | 92% | - |
|  | Alnafisah, K. | Accuracy (multi-label) | 73% | - |
|  | Koyuncu, D. | Area Under Curve | 0.9 | - |
|  | Beamer, G. | Sensitivity | 100% | Positive Predictive Value: 91.8% |
|  | Bug, D. | F1 Score | 83% | - |
|  | Marcos, J. | Probability of Error | 2.93% | - |
|  | Heinemann, F. | Accuracy | 80% | - |
|  | Bartel, A. | Diagnostic Accuracy | 92.3% | Spearman Correlation: 0.98 |
| **Correlation and Scoring Tasks**<br>Studies reporting correlation coefficients and continuous scoring measures | Jule, Y. | Spearman correlation (Ashcroft score) | Dm: $r=0.94$, HDFm: $r=0.96$ | Tissue density: $r=0.83-0.84$, Lung compliance: $r=-0.67-0.71$, FVC: $r=-0.64$ to $-0.68$ |
|  | Murgo, A. | Pearson correlation (Ashcroft score and AI) | 0.86 | - |
|  | Bartel, A. | Spearman's correlation (THS vs Ground Truth) | 0.98 | - |
|  | Heinemann, F. | Pearson's coefficient | 0.92 | - |
|  | Maglogiannis, I. | Mean Absolute error | 0.72 – 0.92 | - |
| **Segmentation and Localization Tasks**<br>Studies reporting spatial accuracy metrics (Dice, mAP, Hausdorff distance) | Harmon, S. | Dice similarity coefficient | 0.873 | Sensitivity: >90% |
| | Salsabili, S. | Dice Similarity Coefficient | 98.22% | Mean Accuracy: 98.34%, Hausdorff distance: 109.68 $\mu\text{m}$, Precision: 93.37%, Recall: 83.47%, F1: 87.87% |
|  | Marzahl, C. (Study 1) | Mean Average Precision (mAP) | 0.90 overall | Equine: 0.88, Human: 0.90, Feline: 0.91, Cross-species: 0.8-0.88 |
|  | Marzahl, C. (Study 2) | Mean Average Precision (mAP) | 0.66 | - |
|  | Bouzin, C. | Percentage Detection in ROI | 75% | - |
| | Vincenzi, E. | Accuracy ($\mu\text{CT}$-histology matching) | 95% | - |
| **Inter-rater Reliability and Concordance Tasks**<br>Studies reporting agreement measures, concordance, and reliability coefficients | Dietert, K. | Concordance (GENIE) | 99.70% | v9 nc algorithm: 99.03%, Overall Accuracy: $\geq 95\%$ |
| | Ben-Hur, A. | Percent Agreement (with LIRA) | 94% | Without LIRA: 87%, Krippendorff's $\alpha$: 0.910 vs 0.796, Error vs expert: 12.1% |
|  | Flores E. | Cohen's Kappa | 0.782 | - |
|  | Salsabili, S. | Fleiss Kappa | 0.84 | - |
|  | Marzahl, C. (Study 2) | Concordance | 0.85 |  |
| **Studies with Incomplete Performance Data** | Ducote, T. | N/A | - | No performance metrics reported |
|  | Palmarini, M. | N/A | - | No performance metrics reported |
|  | Acharya, V. | F1 score | - | Value not specified |
|  | Martinot, A. | Manual inspection | - | Method specified but no quantitative metrics |

Accuracy was the most frequently reported metric across (n=12, 41%), particularly in classification and binary detection tasks, with seven studies reporting values exceeding 90% (Table 2, Supplementary Table S3). Four studies included comprehensive diagnostic metrics beyond accuracy. Segmentation tasks showed robust performance, with Dice similarity coefficients typically above 0.85 and reaching up to 0.98 in select models. Additional spatial and probabilistic metrics were reported less frequently, including Hausdorff distance (30), area under the curve (31), and mean average precision. (20) For continuous outcomes such as fibrosis grading or iron content estimation, correlation coefficients (Pearson or Spearman) between AI and expert assessments often exceeded 0.90.

Only a subset of studies assessed inter-rater reliability for subjective or semi-structured scoring tasks. These included Cohen’s kappa (Lockhart: 0.782), Fleiss ‘kappa (Salsabili: 0.84), and Krippendorff’s alpha (Ben-Hur: 0.796 without AI, increasing to 0.910 with AI assistance) (30, 32, 33). In the latter, agreement between pathologists improved from 87% to 94% when supported by AI, suggesting potential to reduce interpretive variability.

#### Reported limitations

Across studies (Supplementary Table S5), limitations fell into three categories. First, technical challenges included difficulty detecting fine-grained histological features (n=7, 24%) and variability linked to staining quality (n=4, 14%). In addition, pre-analytic factors such as tissue sectioning, slide thickness, scanner resolution, magnification, and artifacts from tissue processing were also noted. Finally, technical resource burdens included computational costs (n=3) and the need for manual oversight (n=3). Second, data quality and annotation issues encompassed inconsistent class assignment (n=5, 17%), lack of gold-standard annotations (n=3, 10%), variability among annotators, underrepresentation of certain classes, small dataset sizes, and the absence of publicly available datasets or code. Third, generalizability and interpretability limitations included the lack of external validation, limited representativity of single sections for whole-organ disease, the absence of model interpretability features, and potential barriers to applying results in multi-institutional settings.

#### Reproducibility and data sharing

Reproducibility practices varied widely. Twelve studies (41%) referenced a code repository (e.g. GitHub), with six providing accessible and fully documented repositories. Most (n=22) used internal or proprietary datasets, and 9 (31%) shared histological image datasets openly or with limited restrictions. Three studies used public datasets that enable cross-study comparison and external validation. None reported standardized protocols or provided a rationale for their approaches to dataset construction, annotation procedures, or reporting practices.

## DISCUSSION

AI holds transformative potential for preclinical pulmonary disease research, particularly in automating and standardizing histological assessments that have long relied on time-intensive and subjective manual scoring. While conventional methods have been foundational, they are prone to inter-observer variability and limit scalability. In contrast, AI tools offer the ability to rapidly analyze large volumes of imaging data with consistent and reproducible outputs.

The diversity observed across the 29 included studies, in terms of dataset characteristics, disease models, staining protocols, and imaging parameters, reflects the field’s dynamic exploration of AI applications across varied experimental settings. Notably, AI tools demonstrated strong performance in identifying morphologically distinct features like tumor regions and granulomas, particularly when integrated into well-optimized histological workflows that include standardized staining (typically H&E or collagen-specific), consistent sectioning, and uniform magnification settings (10x-40x). Preprocessing strategies such as stain normalization, patch-based tiling, and contrast enhancement were widely adopted prior to model training and contributed to improved model robustness.

Our review also identified several gaps that, if addressed, would enhance AI’s potential in preclinical respiratory histopathology. For instance, while AI excels at recognizing well-defined morphological patterns, more nuanced applications of quantitative or semi-quantitative measurements (e.g., fibrosis grading and inflammatory cell quantification) remain challenging. Performance in these tasks often declines in the absence of robust and large-scale training datasets or standardized ground truths.

Another major issue is the lack of standardized quantification methods or scoring systems; as a result, some studies relied on binary annotations, unsupervised clusters, or direct quantification of tissue features such as collagen density or nuclear area. While these approaches facilitated automated analysis, the absence of a standardized ground truth limits cross-study comparability and interpretability. Inconsistent technical reporting presents another challenge, particularly regarding annotation practices and dataset composition was another challenge. Only a minority of studies detailed the number and qualifications of annotators, and few incorporated formal evaluation metrics or explainability tools such as attention maps or saliency visualizations. In addition, most studies to date have focused on proof-of-concept analyses using carefully curated datasets and internal validation. Expanding efforts to include external validation on independent datasets represents a crucial next step for assessing generalizability and strengthening confidence in broader applicability. (26, 34–36) Finally, the dominance of these “black box” AI models, whose decision-making processes remain opaque, is a potential barrier to practical adoption. Improving model transparency is critical to fostering user trust in AI-derived results.

Encouragingly, the success of clinical digital pathology provides a roadmap for preclinical applications. Clinical AI has thrived through large, well-annotated public datasets such as CAMELYON, PANDA, and ImageNet (32–34), achieving expert-level performance in cancer diagnosis, subtyping, and even molecular profiling. (33–36) These advancements were enabled not by model complexity alone, but by transparent validation pipelines, consensus-based annotation, and structured explainability frameworks. Large-scale landmark multicenter studies such as those by Campanella et al. have further demonstrated how interdisciplinary collaboration and clear performance reporting can drive clinical translation. (27, 37) Indeed, studies have suggested that interpretability tools such as attention maps and feature attribution are essential not only for performance but also for end-user trust and regulatory acceptance. (36, 38) This need for explainability has also been strongly advocated in broader reviews and regulatory guidance, highlighting that interpretable AI is essential for adoption in medical practice. (28, 39–41)

While this review provides a comprehensive synthesis of current AI applications in the histological analysis of preclinical pulmonary disease models, some limitations should be acknowledged. First, only English-language publications were included, which may have excluded relevant non-English research. Second, as a scoping review, we did not assess the methodological quality or risk of bias of included studies. As a result, findings from research of varying rigor were synthesized, potentially limiting the strength of conclusions drawn. Finally, the rapid pace of AI tool development means that more recent advances introduced after our literature search may not yet be captured in this review.

The findings from this review highlight both the opportunities promised and current limitations of AI tools in preclinical histopathology. To advance the field, future research should prioritize methodological standardization, transparent and consistent reporting of protocols and annotations, and the use of large, well-curated datasets. Rigorous validation, including testing on independent external datasets from multiple institutions and disease models, will be essential for ensuring robustness and generalizability. Additionally, improving the explainability and interpretability of AI models will help build trust and support practical adoption in research. By addressing these challenges, AI has the potential to enable more efficient, objective, and scalable histological assessment in preclinical studies.

## APPENDIX 1. PRISMA scoping review reporting checklist

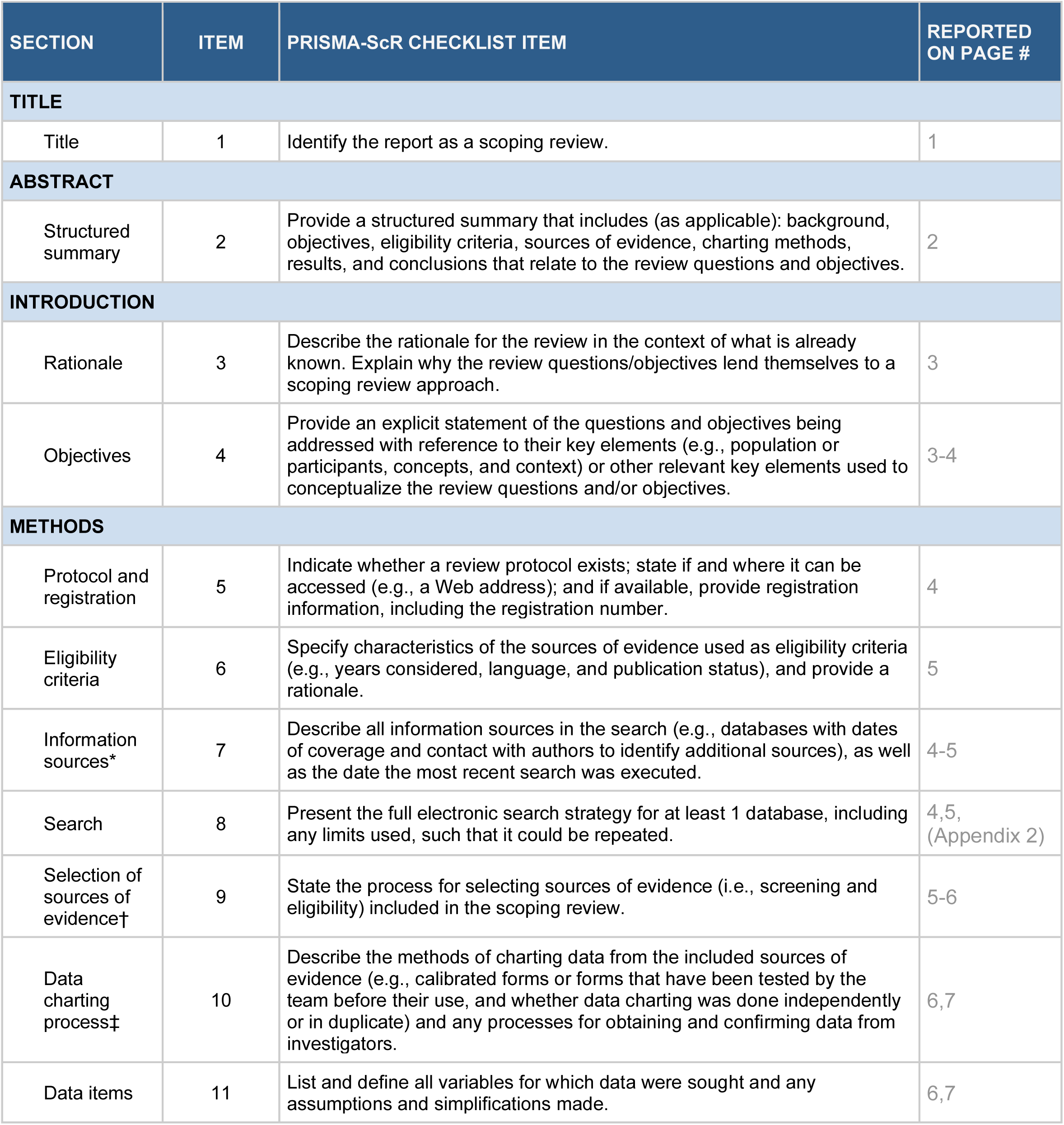
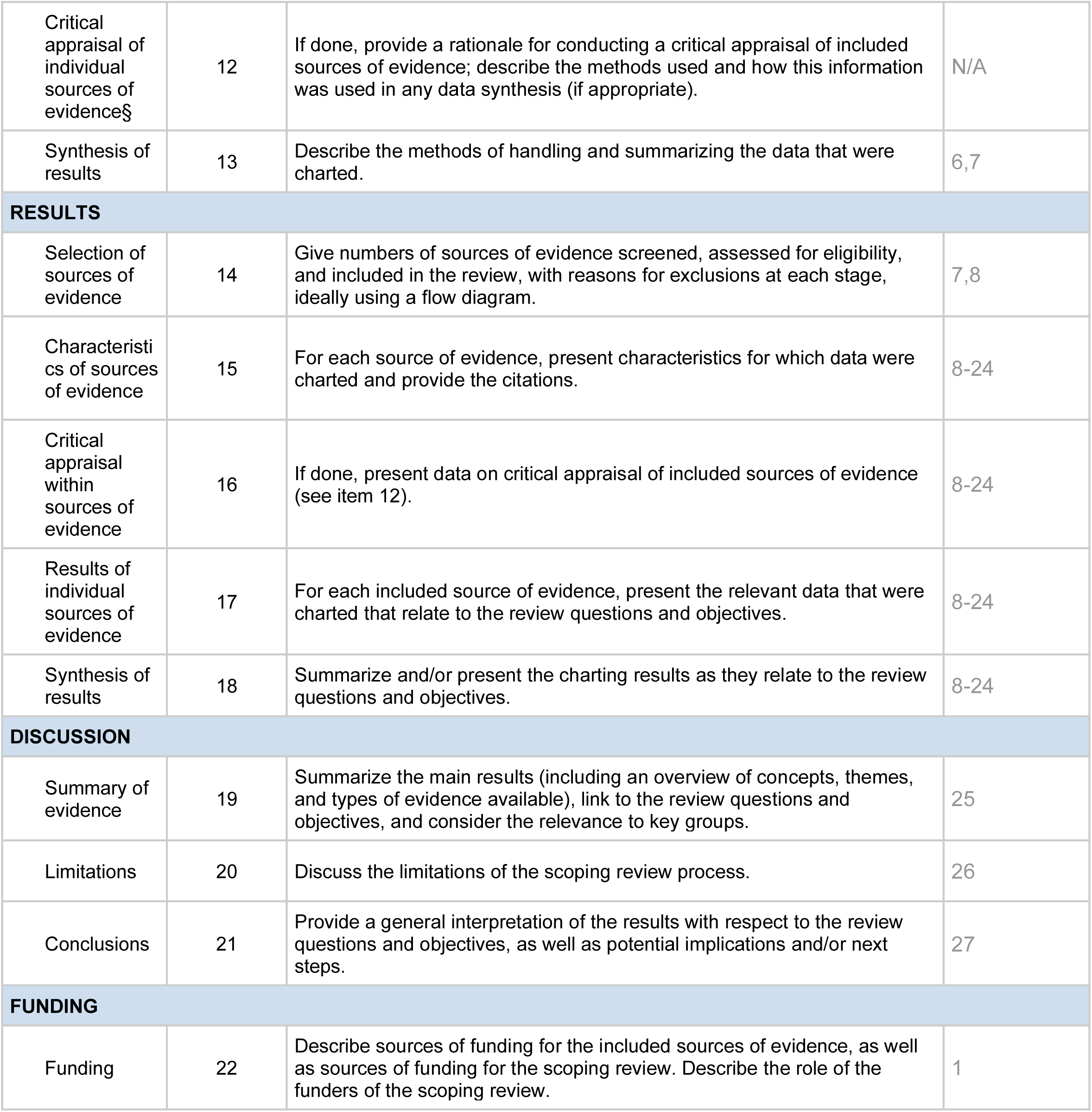

## APPENDIX 2. Search strategy

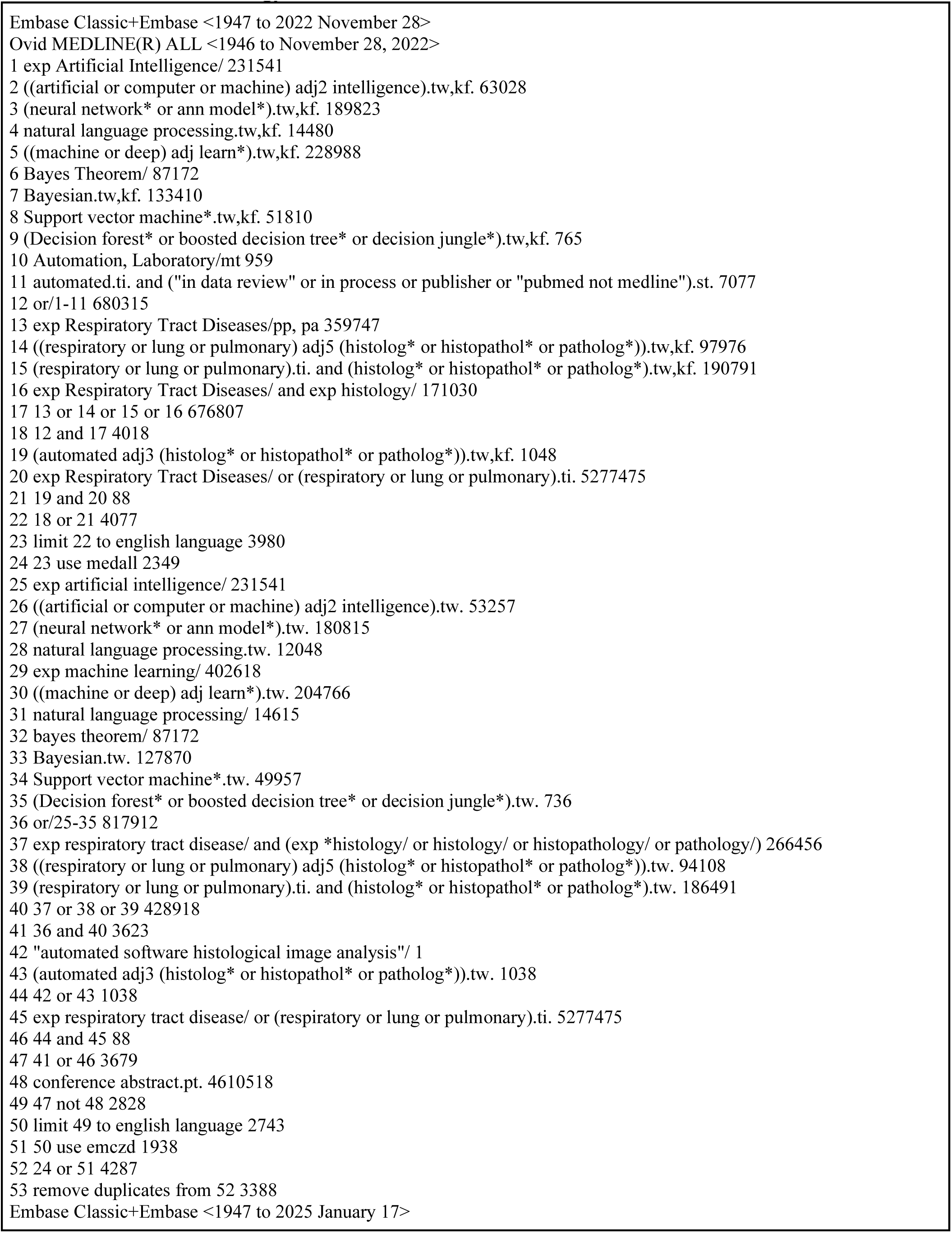
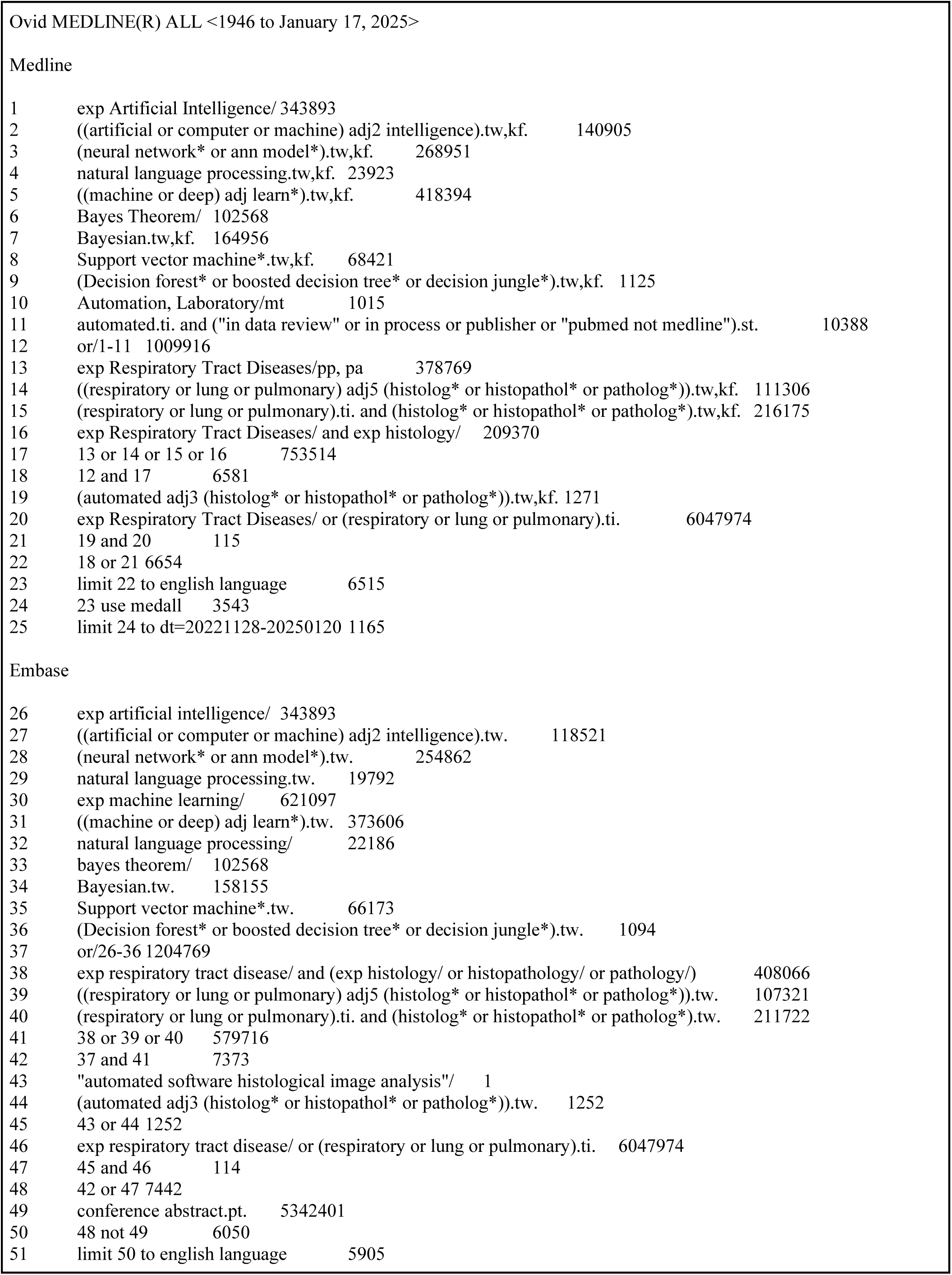
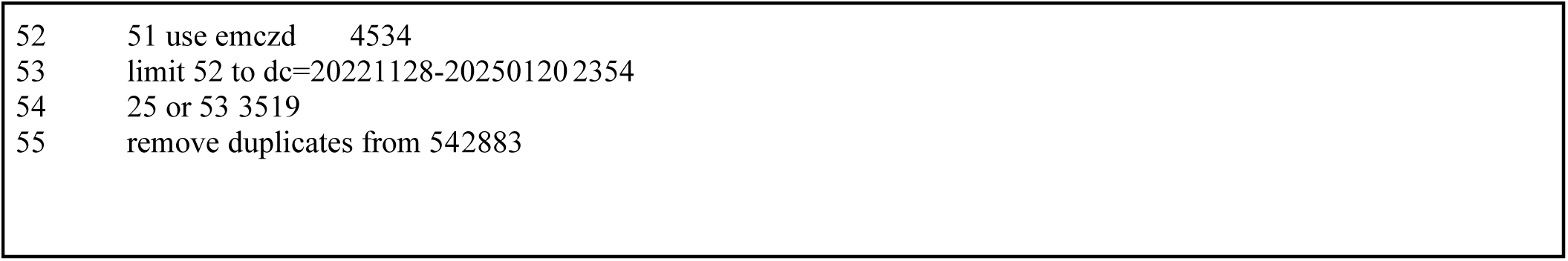

---

**Supplementary Figure S1.**
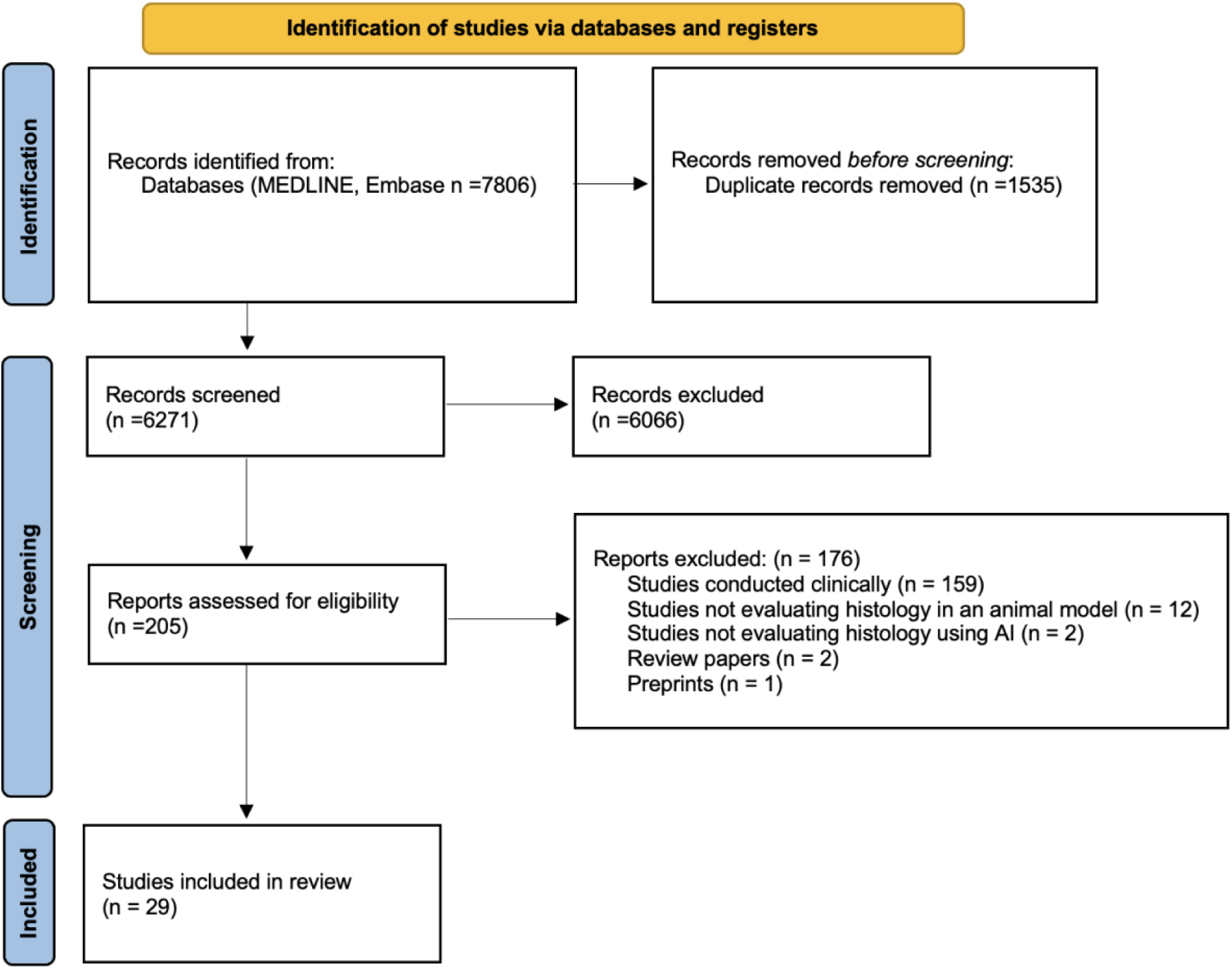
Preferred reporting items in systematic reviews and meta-analysis (PRISMA) flow diagram.

**Supplementary Table S1.** Studies showing the method of staining for slides, the scanoscope used for histological imaging, and the magnification used for the images evaluated.

| **First author** | **Staining method** | **Scanscope used** | **Magnification** |
| --- | --- | --- | --- |
| **Maglogiannis, I** | Hematoxylin and eosin | Nikkon Eclipse E800 (Nikon) | 4x |
| **Marcos, J** | Hematoxylin and eosin | Axioplan 2ie (Zeiss) | 1.25x |
| **Jule, Y** | Masson's trichome | NanoZoomer-SQ (Hamamatsu) | 20x |
| **Dietert, K** | Hematoxylin and eosin | Aperio ScanScope CS2 (Leica) | 400x |
| **Heinemann, F** | Masson trichome | AxioScan Z1 whole slide scanner (Zeiss) | 20x |
| **Seger, S** | Masson's trichome | NanoZoomer 2.0HT whole slide scanner (Hamamatsu) | 40x |
| **Beamer, G** | Hematoxylin and eosin | Aperio ScanScope (Leica) | 400x |
| **Bug, D** | Hematoxylin and eosin | NS | 10x |
| **Ben-Hur, A** | Hematoxylin and eosin | Aperio ScanScope XT (Leica) | 20x |
| **Niazi, M** | Hematoxylin and eosin | Aperio ScanScope (Leica) | 400x |
| **Marzahl, C** | Prussian Blue, Turnbull's blue | Aperio ScanScope CS2 (Leica) | 400x |
| **Bouzin, C** | Picrosirius red | SCN400 slide scanner (Leica) | 20x |
| **Harmon, S** | Hematoxylin and eosin | Aperio ScanScope AT2 (Leica) | 40x |
| **Salsabili, S** | Hematoxylin and eosin | Aperio ScanScope CS2 (Leica) | 20x |
| **Kang, J** | Hematoxylin and eosin | Panoramic Whole-slide scanner 3D (Histotech) | 20x |
| **Marzahl, C** | Prussian Blue, Turnbull's blue | Aperio ScanScope CS2 (Leica) | 400x |
| **Martinot, A** | Hematoxylin and eosin | Not specified | NS |
| **Bartel, A** | Prussian Blue, Turnbull's blue | ScanScope CS2 (Leica) | 400x |
| **Ducote T.** | Hematoxylin and eosin | Aperio Slide Scanner | 20x, 40x |
| **Palmarini M.** | Hematoxylin and eosin | Aperio VERSA 8 Digital Pathology Scanner (Leica Biosystems) | 200x brightfield |
| **Ram S.** | Hematoxylin and eosin | Super COOLSCAN 5000 ED Digital Slide Scanner; Nikon Corporation | 1x objective lens |
| **Pu J.** | Modified Gill's | NS | 20x |
| **Flores E.** | Hematoxylin and eosin | Aperio ScanScope AT2 Slide Scanner (Leica) | 20x |
| **Murgo A.** | Masson's trichome | NanoZoomer S-60 scanner (Hamamatsu) | 20x |
| **Vincenzi E.** | Masson's trichome | Nanozoomer S-60 Digital Slide Scanner (Hamamatsu) | 10x |
| **Liu S.** | Hematoxylin and eosin | ZEISS Axio Scan.Z1, Hohhot, China | 40x |
| **Alnafisah K.** | SHG microscopy | SHG microscopy | NS |
| **Koyuncu D.** | Hematoxylin and eosin | Aperio ScanScope AT2 Scanner / Olympus VS2000 Scanner | 40x |
| **Acharya V.** | Modified Ziehl-Neelsen | NS | 40x |

**Supplementary Table S2.** Summary of the imaging, scoring and AI features of the included studies. Abbreviations: SI, severity index; RBF, radial basis function; WSI, whole-slide images; LIRA, lesion image recognition and analysis; CNN, convolutional neural network; MLI, mean linear intercept; FOV, field of view; SVM, support vector machine; **GLASS-AI,** grading of lung adenocarcinomas with simultaneous segmentation by AI; ISL, iterative selective learning; OPA, ovine pulmonary adenocarcinoma.

| **First author** | **Image resolution** | **Scoring system/parameter for histological classification** | **# of images used to train the AI system** | **Name of the AI program used for automation** | **Are datasets shared?** | **Is the code for AI software publicly available?** |
| --- | --- | --- | --- | --- | --- | --- |
| **Maglogiannis, I** | 0.56 µm/pixel | Interstitial inflammation (0-4), alveolar inflammation (0-3), interstitial fibrosis (0-5) and lung area covered (0-2) | 83,700 training pixels | RBF neural network architecture classifier | No | No |
| **Marcos, J** | 3.54 µm/ pixel | SI of lung area containing emphysematous lesions (normal band, mild band and severe band) | 12 lung sections (6 control mice and 6 severe emphysematous lesion) | Bayesian Framework, large image mosaics used ImageJ | No | No |
| **Dietert, K** | 0.25 µm/ pixel | Quantification of neutrophils, total nuclei and cells, comparison of inflamed lung area and cells density. | Not specified | Aperio Genie histology pattern recognition software for classification. Aperio Nuclear v9 algorithm for quantification | No | No |
| **Heinemann, F** | 0.44 µm/pixel | Ashcroft lung fibrosis score and inflammation score (novel scoring process) | 14,000 labelled tiles for Ashcroft score, ~3,500 images for inflammation score | Inception V3 classifier, scripts developed in Python 3.6 | Yes | Yes |
| **Seger, S** | Not specified | 2 parameters in the Ashcroft lung fibrosis score | 190 WSI | Orbit image analysis software | No | Yes |
| **Ben-Hur, A** | 0.322 µm/pixel | Scoring used seven classification categories: Type I Rim, Type I Core, Type II, Type III, Healthy, Miscellaneous, and Empty slide | 176 digital image scans. Prior to lung scans, 791,000 image patches used for training set | LIRA, scripts developed in Python 3.5 | No | Yes |
| **Harmon, S** | 1 µM/pixel | Binary masks representing 0 (no tumor) and 1 (Tumor) | 137 WSI used for training | Two architectures for segmentation: CNN based on UNet and DeepLabV3+ | Yes | Yes |
| **Salsabili, S** | ~0.5 µm/pixel | MLI score | 14,657 FOV images | CNN based on U-Net architecture | No | Yes |
| **Kang, J** | Not specified | Not specified – identification of tumor lesions | 21,017 images | Inception-v3 deep learning model | Yes | No |
| **Bartel, A** | 0.25 µm/pixel | Total hemosiderin score | 215,426 images | RetinaNet | No | No |
| **Marzahl, C** | 0.25 µm/pixel | 5-tier scoring system of alveolar macrophages based on hemosiderin content | 297,383 images | RetinaNet | Yes | Yes |
| **Martinot, A** | Not specified | 6-feature cumulative scoring system that encompasses the major pathologic features of SARS-CoV-2 disease | Not specified | Random Forest, HALO | No | No |
| **Niazi, M** | 0.23 µm/pixel | The imaging biomarker was quantified as a ratio of necrotic/pyknotic affected regions to non-necrotic tissue | Not specified | Attention Based Pooling, PyTorch | Yes | Yes |
| **Marzahl, C** | 0.25 µm/pixel | A 5-class scoring system for hemosiderophages | 78,047 images | ResNet 18, RetinaNet, support vector machine, regression model | No | Yes |
| **Bouzin, C** | Not specified | Collagen proportionate area in regions of interest and collagen proportion | Not specified | Tissue recognition and segmentation algorithm | No | No |
| **Beamer, G** | Not specified | Granuloma regions are detected, then granuloma regions are searched for cell-poor necrosis | 14 WSI | Linear kernel SVM, Gaussian kernel SVM, Decision Tree, Random Forest, Matlab | No | No |
| **Bug, D** | 2 µm per pixel | Labeled data comprising six distinct tissue classes: tumor (TUM), stroma/connective tissue (MST), necrosis (NEC), blood-cells/-vessels/inbleeding (BLC), vacuoles (VAC) and muscle (MUS) plus an additional class of technical-artifacts (TAR) and background (BGR) class | 71 WSI | Custom algorithm, Fully Convolutional Network, Unet | No | No |
| **Jule, Y** | 3.632 µm/pixel | Two pulmonary tissue density indexes: (i) the mean tissue density (Dm) and (ii) the high tissue density frequency (HDFm) | Not specified | Image analysis, proprietary software program allowing the determination of pulmonary tissue density | No | No |
| **Ducote T.** | 0.5 µm/pixel | Identifying nuclear phenotypes in tumor microenvironment (epithelial, mesenchymal, macrophage, lymphocyte, plasma) | 52 different samples | HALO | No | No |
| **Palmarini M.** | Not specified | Quantification was based on macrophage infiltration (IBA1+ cells), T cell infiltration (CD3+ cells), alveolar epithelial hyperplasia (TTF1+ proliferating pneumocytes) | 30-40 WSI | HALO (DenseNet AI V2 plugin), QuPath | Yes | No |
| **Ram S.** | 0.25 µm/pixel | Automated classification of tumor vs. normal patches | 15 non-tumor images + 15 tumor images (21,934 cancer lesion patches; 653,092 normal patches) | Graph-Based Sparse PCA Network, Classifier: linear SVM | No | No |
| **Murgo A.** | Not specified | Semi-quantitative scoring system was used to assess: edema, hemorrhage, leukocyte infiltration, and epithelial damage. Each scored on a 0–3 scale (0 = absent; 1 = mild; 2 = moderate; 3 = severe) . | Not specified | ilastik | No | No |
| **Vincenzi E.** | Not specified | Ashcroft scoring system (0-8 scale) | Not specified | U-Net | No | No |
| **Flores E.** | 0.502 µm/pixel | Mouse LUAD grading scale (Grades 1–4) | 33 WSIs, ~6000 patches per class across 6 classes | GLASS-AI, Built in MATLAB 2021a, architecture based on ResNet18 with atrous convolution and ASPP layers | Yes | Yes |
| **Chandra D.** | Not specified | Chord length | 1,217 lung images from 137 mice | Deep-Masker, architectures evaluated: U-Net, R2-U-Net, U-Net with ISL, R2-U-Net with ISL | No | No |
| **Alnafisah K.** | Not specified | Quantified collagen content | 49 SHG images from 16 BALB/c female mice | SVM, Random Forest, XGBoost, CNN, LSTM, Autoencoder, plus feature extraction with pre-trained VGG16 | No | Yes |
| **Acharya V.** | Not specified | Classified acid-fast bacilli and activated macrophage nuclei | 18 WSI | Two-layer CNN (Aiforia Cloud v5.1.1) for segmentation, Cell Graph Jumping Knowledge Neural Network for classification, Random Forest, XGBoost, LightGBM, Extra Trees, GraphSAGE, and GATv2 | No | No |
| **Liu S.** | Not specified | Identification and localization of OPA lesions (marked by proliferation of type II alveolar epithelial cells) | 49 WSI, 48,902 lesions | Mask R-CNN | No | No |

**Supplementary Table S3.**
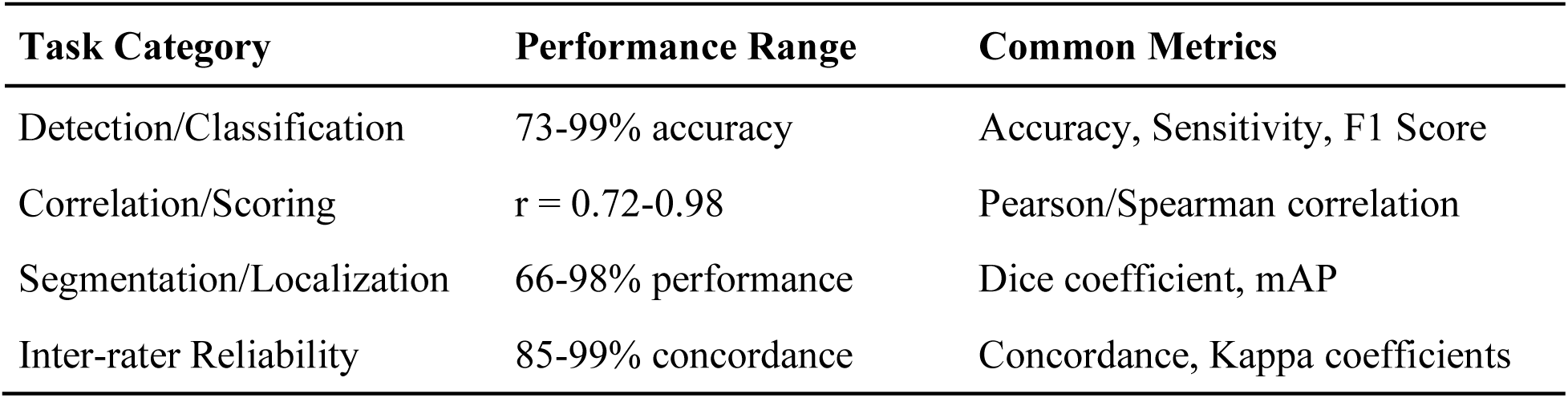
Performance summary by task type. Abbreviations: mAP, mean average precision.

| **Task Category** | **Performance Range** | **Common Metrics** |
| --- | --- | --- |
| Detection/Classification | 73-99% accuracy | Accuracy, Sensitivity, F1 Score |
| Correlation/Scoring | $r = 0.72-0.98$ | Pearson/Spearman correlation |
| Segmentation/Localization | 66-98% performance | Dice coefficient, mAP |
| Inter-rater Reliability | 85-99% concordance | Concordance, Kappa coefficients |

**Supplementary Table S4:**
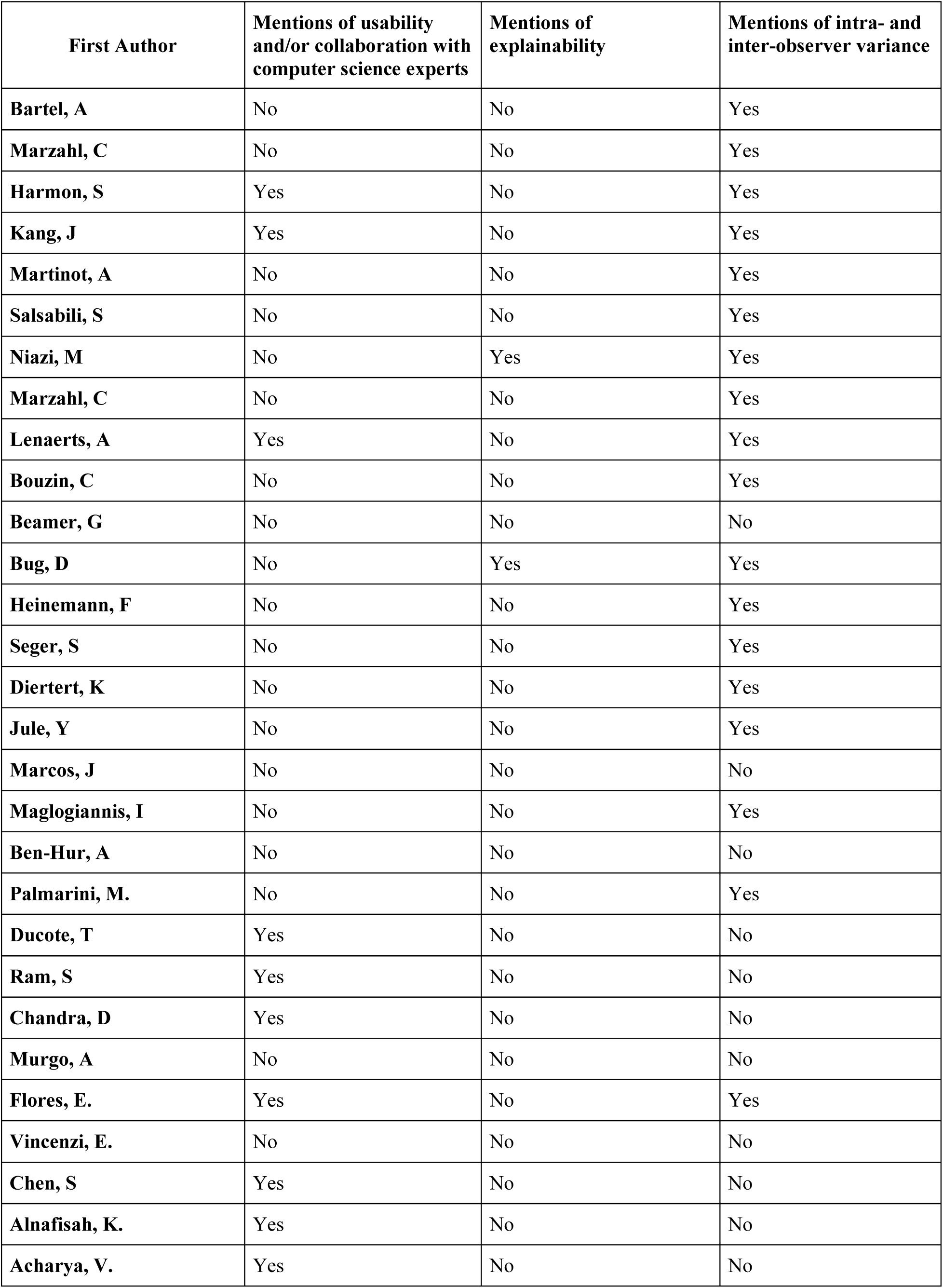
Usability and explainability presented by the included studies.

**Supplementary Table S5.** Artificial intelligence model limitations and key conclusions from included studies.

| **Studies** | **Limitations** | **Conclusions** |
| --- | --- | --- |
| **Maglogiannis, I** | Erroneous predictions on certain histological structures; long training time. | Reduces intra-rater disagreements; highly accurate for fibrosis assessment. |
| **Marcos, J** | Performance depends on tissue preparation and staining; requires human intervention. | Provides a highly accurate tool to assist pathologists. |
| **Jule, Y** | Manual exclusion of large bronchi and vessels may introduce bias; histological scoring methods are observer-dependent and vary across fields, limiting comparability. | Enables precise, automated quantification of bleomycin-induced fibrosis, improving sensitivity at low bleomycin concentrations. |
| **Dietert, K** | Performance depends on tissue preparation and staining; requires human intervention. | Automates immune cell quantification, improving pneumonia analysis in mice. |
| **Heinemann, F** | Excessive training time; ambiguous class assignments; Ashcroft scoring requires expert pathologists and lacks a consensus inflammation score, complicating comparisons. | AI-based Ashcroft scores matched pathologist scores; CNN-based scoring also matched pathologist scores and generates new inflammation metrics without immunohistochemistry, greatly improving efficiency. |
| **Seger, S** | Not specified. | Proposes automated fibrosis analysis, adaptable with further training. |
| **Beamer, G** | Occasionally misclassifies healthy regions as granulomas or necrosis. | Accurately detects granuloma necrosis, aligning with expert pathology. |
| **Bug, D** | Prediction misclassification was observed in about 17% of pixels. | Quantifies biomedical markers and generates interpretable tissue maps for tumor microenvironments. |
| **Ben-Hur, A** | Semi-automated scoring and human intervention is still required. Excessive time is needed for training and assessment | The AI exhibits highly accurate performance. With modification and training, the method can be used on other features. |
| **Niazi, M** | Cannot identify imaging biomarkers of non-supersusceptibility due to class homogeneity. | Enhances interpretability in computational pathology, overcoming deep learning limitations. |
| **Marzahl, C** | No publicly available annotations for feline/human P-Hem slides, limiting transfer learning. | Provides a multi-species P-Hem WSI dataset, supporting inter-species AI development. |
| **Bouzin, C** | Incomplete 3D structure representation; arbitrary outcome thresholds. | PSR staining in brightfield enables accurate collagen fiber detection and fibrosis characterization, with potential for full AI automation. |
| **Harmon, S** | Software usability and user-friendliness issues. | AI segmentation successfully identifies lung adenocarcinoma in mice. |
| **Salsabili, S** | Difficulty in hand-crafting features for blood vessel segmentation; lack of a gold-standard dataset. | A U-Net-based pipeline with color analysis and decision tree classification offers an alternative to manual/semi-automated MLI scoring. |
| **Kang, J** | Erroneous predictions on histological structures; difficulty measuring tumor cells; method is time-consuming and lacks user interactivity. | Inception-v3 detects metastatic tumors in WSI; calls for more user-friendly AI requiring interdisciplinary collaboration. |
| **Marzahl, C** | Single annotator, single lab, and single scanner used for dataset development. | Largest fully annotated EIPH dataset created, supporting deep learning validation and transferability to human samples. |
| **Martinot, A** | Formalin alters immune cell distribution; HALO classifier underperforms in SARS-CoV-2 lung consolidation detection. | Standardized tissue processing enhances cross-institutional vaccine and therapy studies; AI offers alternative disease burden assessments. |
| **Bartel, A** | No gold-standard for unbiased comparison of expert and AI annotations; human annotation errors introduce bias. | AI-based analysis matches human accuracy, scoring thousands of cells in minutes, improving efficiency and reproducibility. |
| **Ducote T.** | HALO AI may misclassify histological features if not trained on diverse patterns, requiring validation. | HALO AI provides cost-effective, reproducible NSCLC tumor microenvironment profiling with strong agreement to immunohistochemistry. |
| **Palmarini M.** | Digital pathology performance varies with staining quality, resolution, and histological differences. | AI-assisted pipeline enables rapid SARS-CoV-2 lung pathology quantification, offering scalable variant virulence assessment. |
| **Ram S.** | Tumor detection methods may produce false positives due to resolution limitations or transitioning cells. | GS-PCANet outperforms existing methods in histopathology image classification with higher accuracy and efficiency. |
| **Pu J.** | Deep-Masker requires extensive training data and struggles with generalizing to different datasets. | Fully automates chord length assessment in murine lung images, reducing variability in COPD research. |
| **Lockhart J.** | GLASS-AI may misclassify normal lung structures and struggles with low-grade tumor distinction. | Efficiently grades lung adenocarcinomas, uncovering tumor heterogeneity with high-resolution analysis. |
| **Murgo A.** | Bleomycin model does not fully replicate human idiopathic pulmonary fibrosis. | Functional readouts like FVC and DFCO enhance its translational potential for anti-fibrotic therapy evaluation. |
| **Vincenzi E.** | Requires manual steps for image alignment and segmentation, limiting full automation. | Effectively integrates histological and μCT data, improving fibrosis characterization and drug discovery. |
| **Chen S.** | Mask R-CNN struggles to differentiate OPA lesions from structurally similar non-lesion areas. | Accurately localizes and diagnoses ovine pulmonary adenocarcinoma, supporting digital pathology transition. |
| **Alnafisah K.** | Small, high-dimensional dataset increases risk of overfitting and limits generalizability without external validation. | Combining statistical and image features enhances lung cancer classification, with SVM performing best. |
| **Koyuncu D.** | AI accuracy depends on high-quality annotations; faces challenges in generalization, class imbalance, and multimodal integration. | Accurately identifies asymptomatic M. tuberculosis infection, outperforming traditional biomarkers. |
| **Acharya V.** | Small dataset limits graph neural network generalizability: model relies on domain-specific graphs and fine-tuning. | CG-JKNN effectively integrates morphology and spatial graphs for tuberculosis detection, with high interpretability. |

**Supplementary Figure S2.**
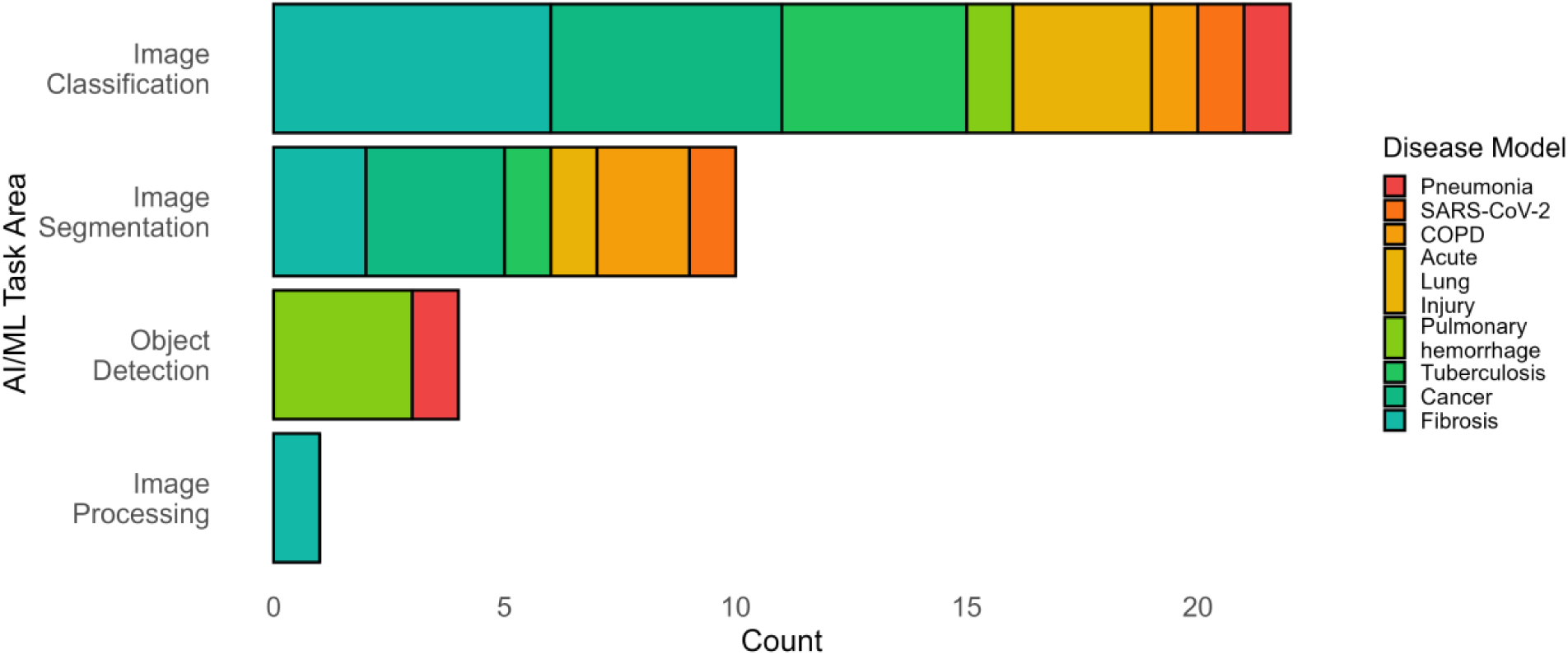
AI/ML Task Areas and Their Associations with Respiratory Disease Models. This bar chart illustrates the frequency of different AI/machine learning (AI/ML) tasks applied in histology studies for respiratory disease models. Each bar is color-coded to represent the associated disease models.

## REFERENCES

1. Meyerholz DK, Beck AP. Principles and approaches for reproducible scoring of tissue stains in research. Lab Invest 98: 844–855, 2018. doi: 10.1038/s41374-018-0057-0.

2. Mazer BL, Homer RJ, Rimm DL. False-positive pathology: improving reproducibility with the next generation of pathologists. Lab Investig J Tech Methods Pathol 99: 1260–1265, 2019. doi: 10.1038/s41374-019-0257-2.

3. Kuo K-H, Leo JM. Optical Versus Virtual Microscope for Medical Education: A Systematic Review. Anat Sci Educ 12: 678–685, 2019. doi: 10.1002/ase.1844.

4. Rizzardi AE, Johnson AT, Vogel RI, Pambuccian SE, Henriksen J, Skubitz AP, Metzger GJ, Schmechel SC. Quantitative comparison of immunohistochemical staining measured by digital image analysis versus pathologist visual scoring. Diagn Pathol 7: 42, 2012. doi: 10.1186/1746-1596-7-42.

5. Uegami W, Bychkov A, Ozasa M, Uehara K, Kataoka K, Johkoh T, Kondoh Y, Sakanashi H, Fukuoka J. MIXTURE of human expertise and deep learning—developing an explainable model for predicting pathological diagnosis and survival in patients with interstitial lung disease. Mod Pathol 35: 1083–1091, 2022. doi: 10.1038/s41379-022-01025-7.

6. Wu B, Moeckel G. Application of digital pathology and machine learning in the liver, kidney and lung diseases. J Pathol Inform 14: 100184, 2023. doi: 10.1016/j.jpi.2022.100184.

7. Birk G, Kästle M, Tilp C, Stierstorfer B, Klee S. Automatization and improvement of μCT analysis for murine lung disease models using a deep learning approach. Respir Res 21: 124, 2020. doi: 10.1186/s12931-020-01370-8.

8. Heinemann F, Birk G, Schoenberger T, Stierstorfer B. Deep neural network based histological scoring of lung fibrosis and inflammation in the mouse model system. PLOS ONE 13: e0202708, 2018. doi: 10.1371/journal.pone.0202708.

9. Deepa BG, Senthil S. Predicting invasive ductal carcinoma tissues in whole slide images of breast Cancer by using convolutional neural network model and multiple classifiers. Multimed Tools Appl 81: 8575–8596, 2022. doi: 10.1007/s11042-022-12114-9.

10. Yu K-H, Wang F, Berry G, Ré C, Altman R, Snyder M, Kohane I. Classifying non-small cell lung cancer types and transcriptomic subtypes using convolutional neural networks. J Am Med Inf Assoc 27: 757–769, 2020. doi: 10.1093/jamia/ocz230.

11. Matute-Bello G, Downey G, Moore BB, Groshong SD, Matthay MA, Slutsky AS, Kuebler WM. An Official American Thoracic Society Workshop Report: Features and Measurements of Experimental Acute Lung Injury in Animals. Am J Respir Cell Mol Biol 44: 725–738, 2011. doi: 10.1165/rcmb.2009-0210ST.

12. Jenkins RG, Moore BB, Chambers RC, Eickelberg O, Königshoff M, Kolb M, Laurent GJ, Nanthakumar CB, Olman MA, Pardo A, Selman M, Sheppard D, Sime PJ, Tager AM, Tatler AL, Thannickal VJ, White ES. An Official American Thoracic Society Workshop Report: Use of Animal Models for the Preclinical Assessment of Potential Therapies for Pulmonary Fibrosis. Am J Respir Cell Mol Biol 56: 667–679, 2017. doi: 10.1165/rcmb.2017-0096ST.

13. Levac D, Colquhoun H, O’Brien KK. Scoping studies: advancing the methodology. Implement Sci IS 5: 69, 2010. doi: 10.1186/1748-5908-5-69.

14. Arksey H, O’Malley L. Scoping studies: towards a methodological framework. Int J Soc Res Methodol 8: 19–32, 2005. doi: 10.1080/1364557032000119616.

15. Peters MDJ, Godfrey C, McInerney P, Munn Z, Tricco AC, Khalil, H. Scoping Reviews (2020). Aromataris E, Lockwood C, Porritt K, Pilla B, Jordan Z, editors. JBI Manual for Evidence Synthesis. JBI; 2024. 10.46658/JBIMES-24-09

16. Russell, Stuart J., and Peter Norvig. Artificial Intelligence: A Modern Approach. 4th ed. Boston: Pearson, 2021. https://aima.cs.berkeley.edu/

17. Mitchell TM. Machine Learning. McGraw-Hill, 1997. ISBN 9780070428072.

18. Dietert K, Nouailles G, Gutbier B, Reppe K, Berger S, Jiang X, Schauer AE, Hocke AC, Herold S, Slevogt H, Witzenrath M, Suttorp N, Gruber AD. Digital Image Analyses on Whole-Lung Slides in Mouse Models of Acute Pneumonia. Am J Respir Cell Mol Biol 58: 440–448, 2018. doi: 10.1165/rcmb.2017-0337MA.

19. Kus P, Gurcan MN, Beamer G. Automatic Detection of Granuloma Necrosis in Pulmonary Tuberculosis Using a Two-Phase Algorithm: 2D-TB. Microorganisms 7: 661, 2019. doi: 10.3390/microorganisms7120661.

20. Marzahl C, Aubreville M, Bertram CA, Stayt J, Jasensky A-K, Bartenschlager F, Fragoso-Garcia M, Barton AK, Elsemann S, Jabari S, Krauth J, Madhu P, Voigt J, Hill J, Klopfleisch R, Maier A. Deep Learning-Based Quantification of Pulmonary Hemosiderophages in Cytology Slides. Sci Rep 10: 9795, 2020. doi: 10.1038/s41598-020-65958-2.

21. Vincenzi E, Buccardi M, Ferrini E, Fantazzini A, Polverini E, Villetti G, Sverzellati N, Aliverti A, Basso C, Pennati F, Stellari FF. A semi-automatic pipeline integrating histological and µCT data in a mouse model of lung fibrosis. J Transl Med 22: 1040, 2024. doi: 10.1186/s12967-024-05819-y.

22. Pu J, Leme AS, De Lima E Silva C, Beeche C, Nyunoya T, Königshoff M, Chandra D. Deep-Masker: A Deep Learning–based Tool to Assess Chord Length from Murine Lung Images. Am J Respir Cell Mol Biol 69: 126–134, 2023. doi: 10.1165/rcmb.2023-0051MA.

23. Maglogiannis I, Sarimveis H, Kiranoudis CT, Chatziioannou AA, Oikonomou N, Aidinis V. Radial Basis Function Neural Networks Classification for the Recognition of Idiopathic Pulmonary Fibrosis in Microscopic Images. IEEE Trans Inf Technol Biomed 12: 42–54, 2008. doi: 10.1109/TITB.2006.888702.

24. Szegedy C, Vanhoucke V, Ioffe S, Shlens J, Wojna Z. Rethinking the Inception architecture for computer vision. In: Proceedings of the IEEE Conference on Computer Vision and Pattern Recognition (CVPR); 2016. p. 2818–26. doi:10.1109/CVPR.2016.308.

25. Ronneberger, O., Fischer, P., Brox, T. U-Net: Convolutional Networks for Biomedical Image Segmentation. n: Navab N, Hornegger J, Wells W, Frangi A, editors. Medical Image Computing and Computer-Assisted Intervention – MICCAI 2015. Lecture Notes in Computer Science. Cham: Springer; 2015. p. 234–41. doi:10.1007/978-3-319-24574-4_28.

26. Chen L-C, Zhu Y, Papandreou G, Schroff F, Adam H. Encoder-Decoder with Atrous Separable Convolution for Semantic Image Segmentation. arXiv preprint arXiv:1802.02611 [cs.CV]. 10.48550/arXiv.1802.02611

27. Acharya V, Choi D, Yener B, Beamer G. Prediction of Tuberculosis From Lung Tissue Images of Diversity Outbred Mice Using Jump Knowledge Based Cell Graph Neural Network. IEEE Access 12: 17164–17194, 2024. doi: 10.1109/ACCESS.2024.3359989.

28. Tavolara TE, Niazi MKK, Ginese M, Piedra-Mora C, Gatti DM, Beamer G, Gurcan MN. Automatic discovery of clinically interpretable imaging biomarkers for *Mycobacterium tuberculosis* supersusceptibility using deep learning. eBioMedicine 62: 103094, 2020. doi: 10.1016/j.ebiom.2020.103094.

29. Bug D, Feuerhake F, Oswald E, Schüler J, Merhof D. Semi-automated analysis of digital whole slides from humanized lung-cancer xenograft models for checkpoint inhibitor response prediction. Oncotarget 10: 4587–4597, 2019. doi: 10.18632/oncotarget.27069.

30. Lithopoulos M, Salsabili S, Sreeraman S, Vadivel A, Thébaud B, Chan ADC, Ukwatta E. Fully automated estimation of the mean linear intercept in histopathology images of mouse lung tissue. J Med Imaging 8, 2021. doi: 10.1117/1.JMI.8.2.027501.

31. Koyuncu D, Tavolara T, Gatti DM, Gower AC, Ginese ML, Kramnik I, Yener B, Sajjad U, Niazi MKK, Gurcan M, Alsharaydeh A, Beamer G. B cells in perivascular and peribronchiolar granuloma-associated lymphoid tissue and B-cell signatures identify asymptomatic *Mycobacterium tuberculosis* lung infection in Diversity Outbred mice. Infect Immun 92: e00263–23, 2024. doi: 10.1128/iai.00263-23.

32. Asay BC, Edwards BB, Andrews J, Ramey ME, Richard JD, Podell BK, Gutiérrez JFM, Frank CB, Magunda F, Robertson GT, Lyons M, Ben-Hur A, Lenaerts AJ. Digital Image Analysis of Heterogeneous Tuberculosis Pulmonary Pathology in Non-Clinical Animal Models using Deep Convolutional Neural Networks. Sci Rep 10: 6047, 2020. doi: 10.1038/s41598-020-62960-6.

33. Lockhart JH, Ackerman HD, Lee K, Abdalah M, Davis AJ, Hackel N, Boyle TA, Saller J, Keske A, Hänggi K, Ruffell B, Stringfield O, Cress WD, Tan AC, Flores ER. Grading of lung adenocarcinomas with simultaneous segmentation by artificial intelligence (GLASS-AI). Npj Precis Oncol 7: 68, 2023. doi: 10.1038/s41698-023-00419-3.

34. Campanella G, Hanna MG, Geneslaw L, Miraflor A, Werneck Krauss Silva V, Busam KJ, Brogi E, Reuter VE, Klimstra DS, Fuchs TJ. Clinical-grade computational pathology using weakly supervised deep learning on whole slide images. Nat Med 25: 1301–1309, 2019. doi: 10.1038/s41591-019-0508-1.

35. Smits AJJ, Kummer JA, de Bruin PC, Bol M, van den Tweel JG, Seldenrijk KA, Willems SM, Offerhaus GJA, de Weger RA, van Diest PJ, Vink A. The estimation of tumor cell percentage for molecular testing by pathologists is not accurate. Mod Pathol Off J U S Can Acad Pathol Inc 27: 168–174, 2014. doi: 10.1038/modpathol.2013.134.

36. Wang X, Janowczyk A, Zhou Y, Thawani R, Fu P, Schalper K, Velcheti V, Madabhushi A. Prediction of recurrence in early stage non-small cell lung cancer using computer extracted nuclear features from digital H&E images. Sci Rep 7: 13543, 2017. doi: 10.1038/s41598-017-13773-7.

37. Singh RV, Agashe SR, Gosavi AV, Sulhyan KR. Interobserver reproducibility of Gleason grading of prostatic adenocarcinoma among general pathologists. Indian J Cancer 48: 488–495, 2011. doi: 10.4103/0019-509X.92277.

38. Nagtegaal ID, Odze RD, Klimstra D, Paradis V, Rugge M, Schirmacher P, Washington KM, Carneiro F, Cree IA, WHO Classification of Tumours Editorial Board. The 2019 WHO classification of tumours of the digestive system. Histopathology 76: 182–188, 2020. doi: 10.1111/his.13975.

39. Amann J, Blasimme A, Vayena E, Frey D, Madai VI, Precise4Q consortium. Explainability for artificial intelligence in healthcare: a multidisciplinary perspective. BMC Med Inform Decis Mak 20: 310, 2020. doi: 10.1186/s12911-020-01332-6.

40. Marey A, Arjmand P, Alerab ADS, Eslami MJ, Saad AM, Sanchez N, Umair M. Explainability, transparency and black box challenges of AI in radiology: impact on patient care in cardiovascular radiology. Egypt J Radiol Nucl Med 55: 183, 2024. doi: 10.1186/s43055-024-01356-2.

41. Ali S, Akhlaq F, Imran AS, Kastrati Z, Daudpota SM, Moosa M. The enlightening role of explainable artificial intelligence in medical & healthcare domains: A systematic literature review. Comput Biol Med 166: 107555, 2023. doi: 10.1016/j.compbiomed.2023.107555

